# Combinatorial perturbation structure enables graded inference of signaling activity by CIRCE

**DOI:** 10.64898/2026.11.19.767979

**Authors:** Nicholas T. Hutchins, Samuel Maffa, Claire Lu, Pulin Li

## Abstract

Cells decode external inputs into intracellular activities of varying magnitude, and inferring both input identity and strength is crucial for cell-fate engineering. However, perturbation-response models typically treat all cells exposed to a given input as a discrete class, ignoring cell-to-cell variation in response magnitude. Here, we introduce a structured-label framework that infers both input identity and strength from combinatorial perturbation screens without dose-resolved training labels. We represent each perturbation combination in a binary combinatorial space and replace one-hot supervision with a Boltzmann distribution that decays with the Hamming distance between combinations. This approach preserves classification accuracy while improving calibration and resolving continuous pathway activities across individual cells. Importantly, intermediate scores primarily reflect ambiguity in cellular response states rather than insufficient model knowledge. Applied to engineered and natural tissues, the calibrated model successfully recovered graded activities. These results establish combinatorial perturbation screens as a self-calibrating framework for decoding graded cellular responses.

## INTRODUCTION

While external inputs drive cell fate changes and physiological adaptation, the exact choice a cell makes depends not only on the identity of the input, but also the magnitude of the response within the cell ^1–4^ . The classic example is tissue patterning in developing embryos, where spatially graded ligand concentrations instruct the formation of adjacent cell-type domains that are distinct from one another ^1^. However, response magnitudes not only depend on the input concentration, but also the internal cellular state. Different cell types respond to the same input with different strengths. Similarly, signaling pathways often act in parallel to alter the internal state of the cell, and thus the response magnitude to one pathway is affected by the presence of other pathways ^5–8^. Furthermore, even clonal populations of cells exposed to identical inputs can respond with different magnitudes due to noise, leading to distinct fate choices ^9–11^. Therefore, resolving the magnitude of signaling response, by taking into consideration of both the external input and internal cell state, is essential for understanding how cells make decisions and for engineering cell fates towards therapeutic intervention and regeneration medicine^12^.

Many machine learning models trained on perturbation data do not explicitly model perturbation magnitude. Instead, they frame perturbation prediction as a binary classification task: determining whether a perturbation is present or absent ^13–18^. This formulation is inherited from cell-type classification objectives, which are intrinsically binary ^19–21^. Consequently, it masks the cell-to-cell variation in response magnitude within the training data. These models therefore have limited capability to recover the response magnitude in target cell types, which can vary substantially with biological context. They may also perform poorly when a response magnitude lies in an intermediate state between the nominal perturbed and unperturbed classes ^22^. Directly training such models to infer perturbation strength would require perturbation data spanning many precisely measured doses for each perturbation to create hard dose labels, which remain rare ^23^. Moreover, this strategy would become impractical as the number of perturbed pathways increase, particularly when combinations of perturbations and their doses must both be considered.

Perturbation screens, often performed as combinatorial inputs, however, contain a largely unused source of information about response magnitude. When a pathway is activated in different combinations, each combination places that pathway in a distinct cellular context ^12,24^. Crosstalk among pathways can consequently alter the transcriptional response, generating a range of effective response magnitudes even when the nominal label for each pathway is binary. Applying the same perturbation across diverse starting cell states can further broaden this distribution. Thus, the combinatorial structure of perturbation screens may inherently sample a range of signaling response magnitudes, including intermediate states, that can be used to calibrate graded model predictions.

Here, we develop a structured-label framework for graded signaling inference from combinatorial perturbation screens. We replace one-hot multiclass targets with a distribution defined by Hamming distance in the combinatorial label space, and implement this objective in a scANVI-based model that we term Calibrated Intracellular Response Continuum Estimation (CIRCE). CIRCE improves calibration without sacrificing categorical performance. Consequently, it yields substantially more intermediate pathway scores than binary or conventional multiclass models, and these intermediate predictions reflect true biological response heterogeneity. Finally, CIRCE recovers known graded signaling in engineered and native tissues with tight single-cell response predictions across spatial positions. Thus, combinatorial perturbation screens can be used to calibrate a continuous representation of signaling activity from ostensibly binary labels.

### Combinatorial structure of perturbation screens provides an internal scale for graded signaling inference

We sought to leverage the combinatorial structure of perturbation screens to infer the relative activation strength of defined signaling pathways from single-cell transcriptional states **(Fig. 1a, *left*)**. When a pathway is activated alone and in several co-stimulation conditions, its nominal label remains “on” in every condition. However, interactions with co-active pathways can alter both the magnitude and composition of its transcriptional response. Across cells and perturbation combinations, the screen therefore samples a distribution of response magnitudes for each pathway. We hypothesized that this combinatorial structure could provide an internal calibration scale for converting binary perturbation labels into graded pathway activity scores.

**Figure 1.**
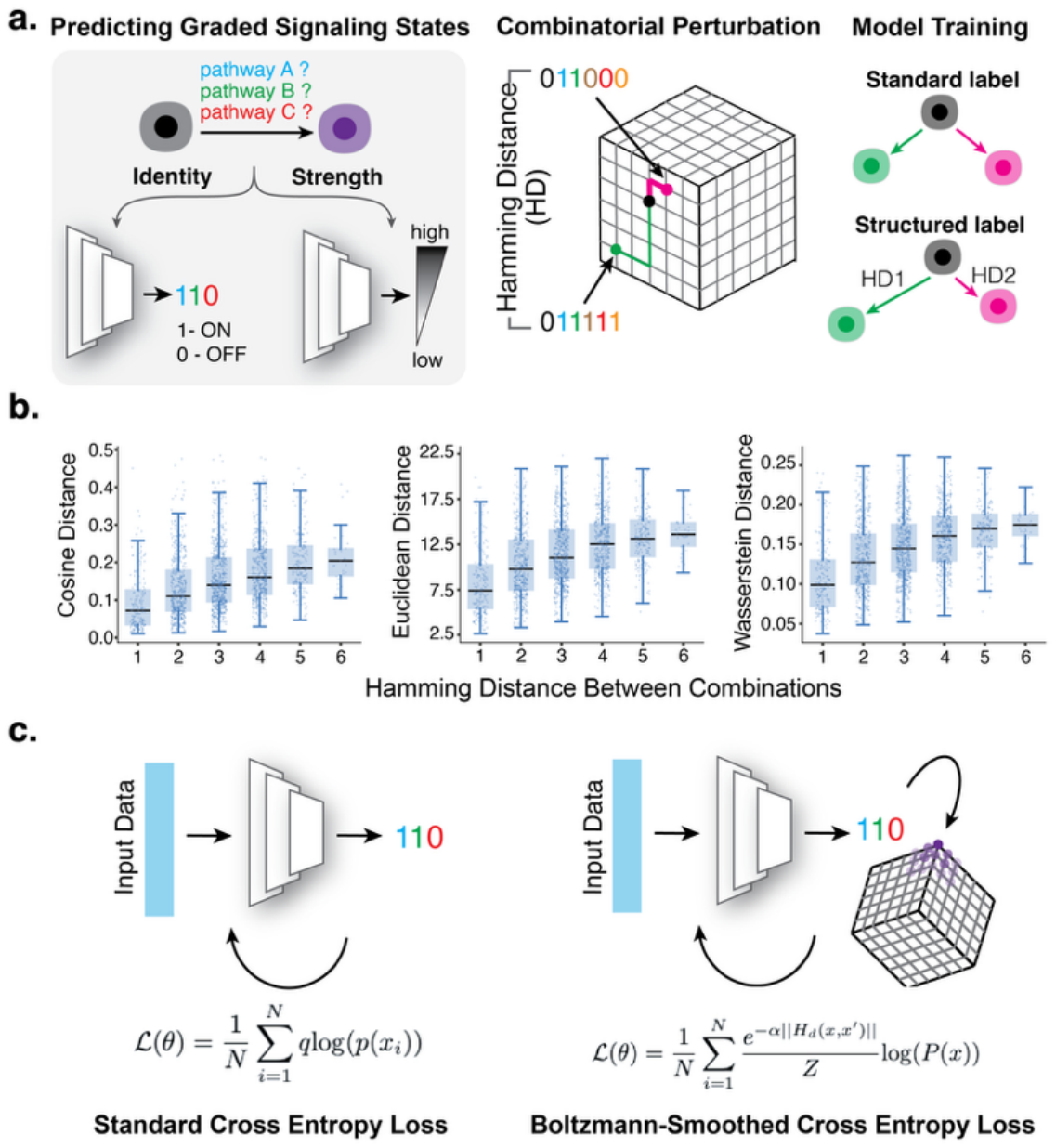
Combinatorial structure of perturbation screens provides an internal scale for graded signaling inference. **a**, Signaling perturbations vary in pathway identity, combination, and effective response strength. *Left*, the goal of signaling inference is to identify both the activation state and strength of the signaling pathway in individual cells. *Middle*, schematic of a multidimensional combinatorial signaling space, in which Hamming distance between combinatorial conditions quantifies the dissimilarity between the signaling environments of two cells. *Right*, models may be trained using objectives that ignore this combinatorial structure or using modified objectives that incorporate distances among perturbation combinations. **b**, Distance between combinatorial signaling labels positively correlates with the distance between their transcriptomic profiles (Spearman’s *ρ*: Cosine 0.371, p=1e-64; *ρ*: Euclidean 0.359, p=2e-60, *ρ*: Wasserstein 0.369, p=4e-64). Cell pairs separated by defined Hamming distances were sampled, and transcriptomic dissimilarity was quantified using three distance metrics (cosine, Euclidean, and Wasserstein). **c**, Comparing different ways of calculating cross entropy loss. Standard cross-entropy assigns all target probability to the observed perturbation combination. In contrast, the Hamming-Boltzmann objective distributes target probability across combinations according to their Hamming distance from the observed state, assigning greater weight to neighboring signaling states.

To determine whether combinatorial labels have a defined geometry that reflects transcriptional response similarity, we analyzed combinatorial signaling perturbation screens in mouse and human pluripotent stem cells ^13,25^ . Stem cells were treated with diverse, randomly selected ligand combinations, and the transcriptional states of the resulting differentiated cells were profiled by single-cell RNA-sequencing (scRNA-seq). Importantly, each ligand was applied at a single fixed concentration across all combinations. To encode this combinatorial label structure, we represented each combination of the six pathways (BMP, FGF, RA, SHH, TGF-β, and WNT) as a six-bit vector, and the geometric distance between two combinations as the Hamming distance, which is the minimum number of bit flips required to interchange two binary sequences **(Fig. 1a, *middle*)**. For pairs of cells, we compared the distance between their combinatorial signaling labels with the distance between their transcriptomic profiles. Cells exposed to more similar signaling combinations also exhibited more similar transcriptomic profiles **(Fig. 1b)**. To confirm that this relationship was not confounded by the high dimensionality of the data, we independently evaluated it using shared-nearest-neighbor connectivity and observed the same association **(Supplementary Fig. 1)** ^26^. Together, these analyses support the idea that the combinatorial perturbation structure encodes information about the relative strength of pathway activation.

### Incorporating combinatorial geometry in model training

We reasoned that a model that preserves geometric relationships among perturbation combinations could use these response states to learn graded pathway scores, even when the experimental labels remain binary **(Fig. 1a, *right*)**. Existing label-classification approaches generally fall into two categories. Standard binary classifiers treat each pathway as independent and derive a combinatorial prediction by concatenating individual pathway-specific output, whereas conventional multiclass classifiers treat each pathway combination as a separate category ^27^. Neither approach leverages the inherent combinatorial geometry of the data to represent the relationships among nearby combinations in label space. By encoding this geometry during training, a model should be encouraged to distribute probability across biologically neighboring combinations and, after marginalization, produce graded scores for individual pathways.

To incorporate the geometric relationships among perturbation combinations, we changed how model is taught what constitutes a correct prediction. In conventional model training, each cell is assigned a one-hot target, and the model is trained with standard cross-entropy loss that treats all alternative combinations equally incorrect. In contrast, we proposed a structured approach using Boltzmann-smoothed cross-entropy loss, which assigns the highest probability to the observed combination but gives progressively lower probability to combinations that are farther away in Hamming distance **(Fig. 1c)**. Specifically, for a cell with observed combination y, we replaced the one-hot target with a soft target distribution over all combinations *c*_*j*_:

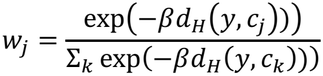

Where *d*_*H*_ is the Hamming distance between combinations, and β controls how strongly probability is concentrated around the observed combination. Optimizing cross-entropy against this structured target assigns partial credit to predictions that correctly match the observed condition for some, but not all, pathways. This objective should enable the model to learn representations that reflect both combinatorial label structure and the graded transcriptional response states associated with it.

We implemented this objective in a shared scVI/scANVI backbone ^19,28^. In selected models, we annealed β from low to high during training, allowing the model to first learn the global geometry of the label space before resolving individual combinations more sharply. In other models, β was held constant. For comparison, we also constructed four other training objectives that incorporated multiclass geometry in distinct ways: independent binary classifiers (“Binary”), a conventional 64-class classifier (“Multiclass”), a multitask model with six binary heads (“MultiTask”), and a joint model that combines binary and multiclass objectives (“Disentangled”). For each model, we derived pathway-specific activity scores by marginalizing the predicted probability over all combinations that contained the pathway of interest. This common readout enabled direct comparison of categorical performance, probability calibration, and the prevalence of graded pathway scores across models.

### Structured-label smoothing generates more graded prediction through improved calibration

We first evaluated whether the inferred pathway score by each model supported a graded probabilistic interpretation. For this to be true, predicted probabilities must correspond consistently to observed outcome frequencies. For example, among cell-pathway pairs assigned an activation probability of 0.8, the pathway should be active in approximately 80% of cases. This property, known as model calibration, can be quantified by the expected calibration error (ECE), which measures the discrepancy between predicted probabilities and observed outcome frequencies across probability intervals **(Fig. 2a)** ^29^.

**Figure 2.**
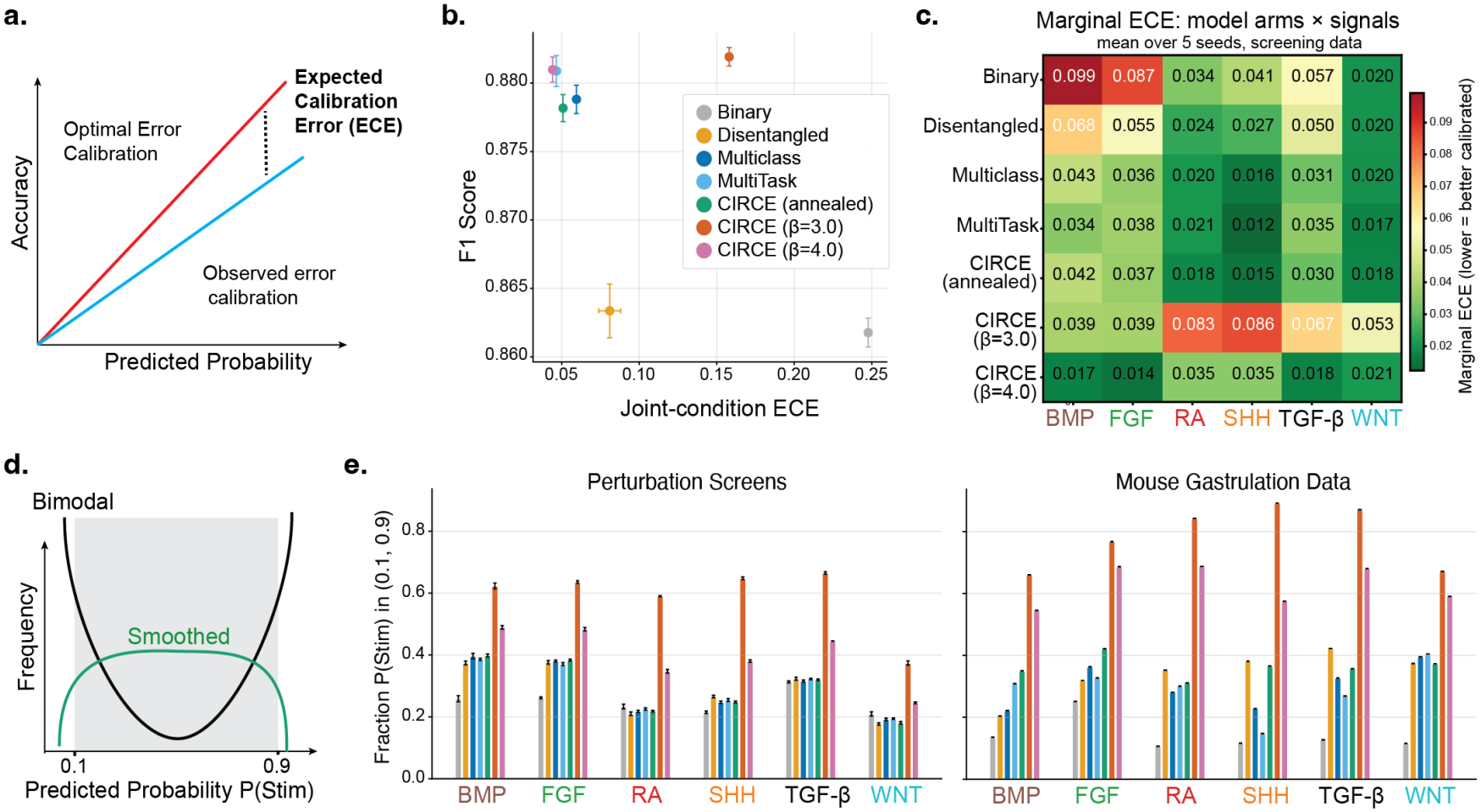
Structured-label smoothing improves calibration and produces more graded pathway scores. **a**, Expected calibration error (ECE) quantifies the discrepancy between predicted probabilities and observed outcome frequencies across probability intervals. Under perfect ECE, the observed frequency of true positives equals the predicted probability across the full range of predicted values. **b**, Joint condition ECE measures the calibration error across all possible multiclass combinations. Joint condition ECE is plotted against model accuracy, measured by F1 score. **c**, ECE after marginalizing combination-level probabilities to individual pathway probabilities. **d**, Improved error calibration (*green*) shifts distribution of predicted probabilities toward an intermediate range, releative to the bimodal distribution produced without calibration (*black*). Stim denotes the activation label for a given pathway, where 0 indicates an inactive state and 1 indicates an active state. **e**, Fraction of predictions within the intermediate pathway score range, defined as probabilities between 0.1 and 0.9, as illustrated in **d**. *Left*, predictions for the human and mouse stem cell perturbation screens ^13,25^; *right*, prediction for an independent mouse gastrulation atlas ^42^. Colors denote distinct models, using the same color scheme in **b**. Error bars indicate the standard error of the mean across held-out cells from the screens (n=7,659; *left*) or all cells (n=130,000; *right*) in the gastrulation atlas.

We evaluated model calibration using the same combinatorial signal screens on mouse and human pluripotent stem cells ^13,25^. These datasets were collected as five batches, each representing distinct differentiation lineages (organ-specific mesenchyme, endoderm, primitive streak, and early mesenchyme). In the train-test split, we held out one batch and its associated cell types, trained each model on the remaining batches, and evaluated on the held-out cell types. Across all train-test splits, models incorporating joint combination information were better calibrated than independent binary models. CIRCE (with β =4.0) achieved the lowest overall ECE and maintained high accuracy **(Fig. 2b)**. The CIRCE (β =3.0) model has low accuracy and high ECE which may be the result of generally high uncertainty as the training loss does not emphasize the correct combination as strongly. The CIRCE annealed model has properties similar to the multiclass model because the loss converged to the standard multiclass loss during training and it does not benefit from the combinatorial structure of the signaling.

Improved calibration was also evident after marginalizing predictions to individual pathways: CIRCE and the other multiclass models reduced pathway-level ECE across all six pathways, with CIRCE being the most pronounced on average **(Fig. 2c, Supplementary Fig. 2)**. Thus, CIRCE outperformed other models in improving calibration, which did not arise from reduced classification performance; rather, structured smoothing improved the quality of the predicted probabilities.

To further compare the capability of generating graded predictions among models, we quantified how frequently each model assigned intermediate pathway scores, defined as values between 0.1 and 0.9 **(Fig. 2d)**. When tested on both the perturbation screens and an independent mouse gastrulation atlas ^42^, the Binary independent and conventional Multiclass models concentrated predictions near zero or one, whereas CIRCE produced a substantially larger fraction of intermediate scores **(Fig. 2e)**. These results indicate that structured supervision uses diversity of responses across combinations to learn a more graded representation of each pathway.

Furthermore, we asked whether these gains could be achieved by generic label smoothing alone. We applied conventional label smoothing to the Binary model by decreasing the target probability by a fixed value (ε) and relocating that probability to the alternative class during cross-entropy training. We found that this improved calibration over a broad range of smoothing strengths without materially reducing exact-match accuracy, indicating that the original binary model was overconfident **(Supplementary Fig. 3)**. However, CIRCE further improved calibration by distributing target probability according to the known geometry of the combinatorial perturbation space, rather than treating all alternative labels as equally plausible. Thus, CIRCE prevents over confidence and learns a more graded representation of probabilities.

Finally, we examined how model performance varied with the number of cells sampled per signaling combination. Predictive performance plateaued at approximately 80 cells per combination, suggesting that, beyond this point, expanding coverage of distinct combinations is more valuable than collecting additional cells from already sampled condition **(Supplementary Fig. 4)**.

### Intermediate pathway scores predominantly reflect variation in cellular responses

Next, we asked whether intermediate pathway scores reflect genuine variation in cellular response magnitude. An intermediate prediction can arise for two distinct reasons. First, it may reflect epistemic uncertainty, because the model has not observed sufficiently similar transcriptional states during training and therefore lacks confidence in its prediction. Alternatively, it may reflect aleatoric ambiguity, because the cell occupies a response state that does not fall definitively into an individual perturbation class **(Fig. 3a)**. The latter possibility is consistent with the biological premise of our model, in which hard treatment labels may conceal a continuous distribution of cellular response magnitudes to the same nominal perturbation.

**Figure 3.**
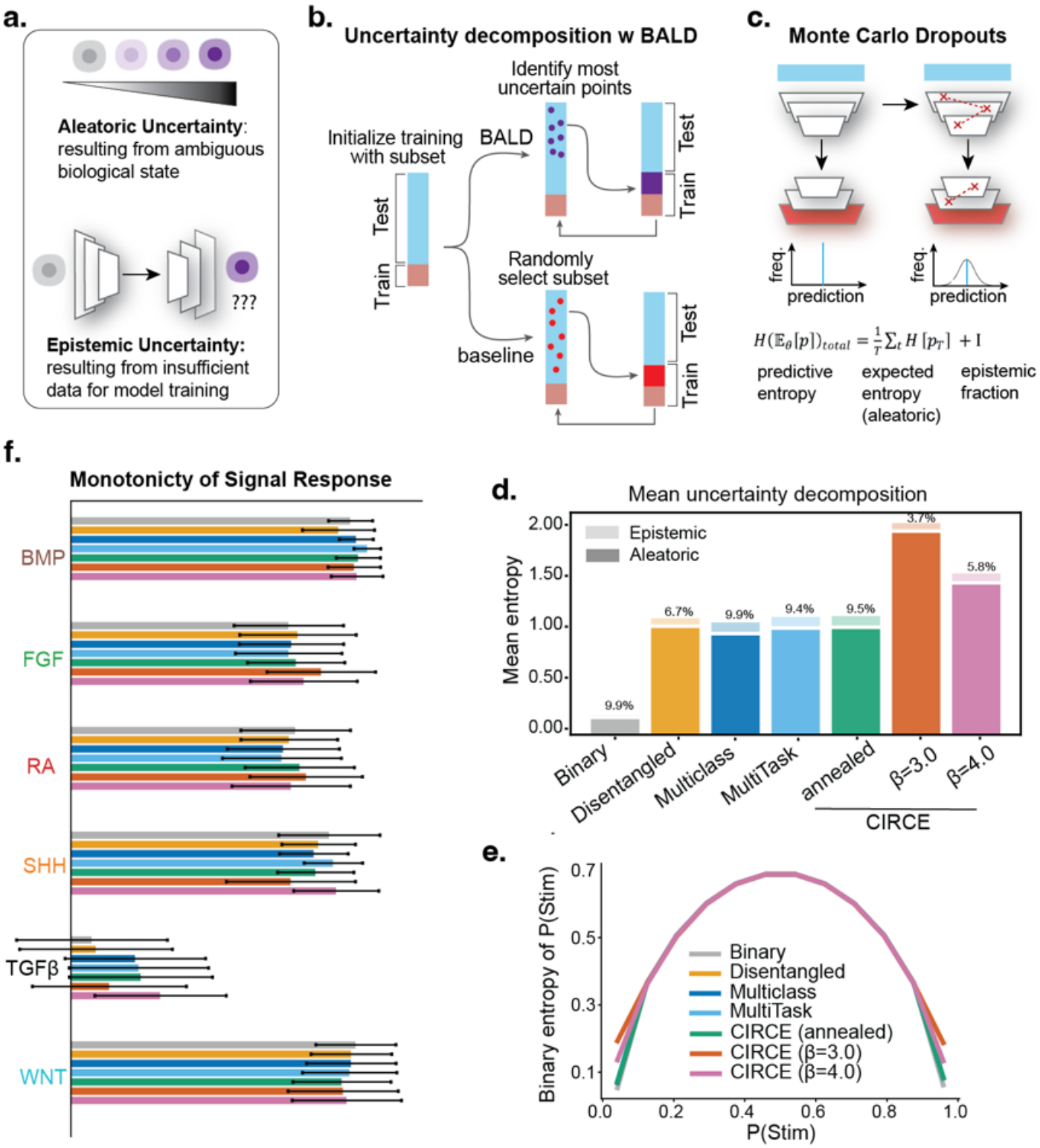
Intermediate pathway scores reflect heterogeneity in cellular response state. **a**, Different sources of uncertainty in model prediction. **b**, An active-learning framework enables uncertainty decomposition. The framework withholds subsets of cells from the training data, quantifies uncertainty of each held-out prediction, and iteratively adds selected cells back to the training set. Bayesian Active Learning by Disagreement (BALD) selects cells with the highest uncertainty scores, whereas the baseline model randomly selects subsets of cells. **c**, Quantifying predictive uncertainty with Monte Carlo (MC) dropout. MC dropout estimates the distribution of outputs by enabling dropouts at inference time. Uncertainty can be decomposed into aleatoric and epistemic components the difference between total predictive entropy and expected entropy. **d**, Decomposition of epistemic and aleatoric uncertainty by BALD across all models, using stem cell screening data. **e**, Relationship between total predicted uncertainty and predicted pathway activation score P(Stim). Total predicted uncertainty is computed from the entropy of the output distribution from MC dropout. **f**, Correlation between graded pathway scores with intermediate levels of pathway activation, measured by monotonicity. Pathway activation was quantified based on the expression levels of canonical pathway-response genes, normalized against null distribution of a randomly chosen gene set.

We used a Bayesian Active Learning by Disagreement (BALD) approach to distinguish these possibilities ^30–33^. We first withheld subsets of cells from the training data, quantified uncertainty of each held-out prediction, and iteratively added selected cells back to the training set before retraining the classifier **(Fig. 3b)**. If uncertainty primarily reflects missing coverage of signaling state space, acquiring the most uncertain cells should improve performance more rapidly than random acquisition. In contrast, if uncertainty primarily reflects intermediate biological states between two perturbation classes, adding uncertain cells should provide little advantage over random sampling.

We quantified uncertainty using Monte Carlo dropout (MC dropout), which generates multiple predictions for each cell **(Fig. 3c)** ^31^. Total uncertainty was quantified as predictive entropy, which measures the spread of probabilities assigned to each of the classes ^34^. As expected, predictive entropy increased as performance declined across all models and signaling pathways, confirming that this metric identified difficult or ambiguous predictions **(Supplementary Fig. 5)**.

We further decomposed total uncertainty into an aleatoric component, calculated as the entropy of the forward passes through the network, and an epistemic component, calculated as the difference between total entropy and aleatoric uncertainty. We validated this decomposition using synthetic datasets with known sources of uncertainty. These synthetic data sets were generated with a 40-gene response indicator block which is increased in the positive signaling cases along with 200 non-response genes. To create an aleatoric-dominated case, we introduced label noise by flipping individual pathway labels. To create an epistemic-dominated setting, a synthetic gene set composed of 120 genes with random expression values was simulated in cells. This represented unobserved gene expression for the model. In each round of training, the model was introduced to these cells and can learn the synthetic gene expression. In both cases, the estimated uncertainty decomposition correctly recovered the known dominant source of uncertainty **(Supplementary Fig. 6)**.

We then applied this framework to evaluate uncertainties of model predictions on the screen datasets. Across model classes and across both in-sample and held-out-batch evaluations, BALD-guided acquisition did not outperform random sampling, measured by either exact-match accuracy or Hamming distance **(Supplementary Fig. 7)**. In several settings, random acquisition performed better than BALD. Consistent with this result, uncertainty decomposition showed that aleatoric uncertainty dominated across all models **(Fig. 3d)**. At the single-cell level, intermediate pathway scores were associated with higher estimated aleatoric uncertainty and disagreement with the binary treatment labels. (**Fig. 3e; Supplementary Figs. 8-9)**. These results indicate that ambiguous predictions arise predominantly from intermediate biological states between two perturbation classes, rather than from insufficient training coverage, and thus intermediate pathway activation scores might represent graded pathway activities.

To further evaluate whether these intermediate pathway activation scores align with graded activation of signaling pathways, we measured their correlation with the levels of canonical signaling response genes **(Table 1)** ^35^. The correlation was normalized to a null distribution of randomly chosen genes and quantified as the monotonicity. Canonical response genes varied continuously with the corresponding pathway activation scores, confirming that the output probabilities align with the strength of signaling response **(Fig. 3f)**. Together, these analyses support the interpretation that intermediate scores capture real heterogeneity in how strongly cells respond to a nominal perturbation, rather than simply identifying regions in which the model lacks training data.

**Table 1.** Context Independent Response Gene Sets Across Pathways.

| **Pathway** | **Response Genes** |
|---|---|
| **RA** | *CYP26A1, HOXB1, HNF1B, CYP26C1, HOXA1, HOXB2, HOXA3, HOXA2, HOXB3* |
| **BMP** | *ID2, BMPER, ID4, ID1, BAMBI, MSX1, ID3, MSX2, SMAD7* |
| **FGF** | *SPRY1, SPRY2, DUSP6, SPRY4, ETV5, ETV4, FOS, MYC, JUNB, DUSP14* |
| **WNT** | *AXIN1, CCND1, DKK1, MYC, NOTUM, AXIN2, SP5, LEF1* |
| **TGF-β** | *SMAD7, TWIST1, TWIST2* |
| **SHH** | *GLI1, PTCH1, HHIP, FOXF1, FOXF2* |

### Graded scores recover imposed signaling gradients in a controlled *in vitro* system

To experimentally test whether the graded pathway scores corresponded to independently defined differences in signaling strength, we analyzed a human endoderm patterning system in which morphogen gradients were imposed through a microfluidic system^36^. BMP, WNT, and FGF were provided from the posterior to form posterior-to-anterior concentration gradients, whereas anterior identity was maintained by BMP and TGF-β inhibition **(Fig. 4a)**. This dataset provides a stringent test because the model was trained on discrete combinations but evaluated on cells exposed to continuous spatial gradients.

**Figure 4.**
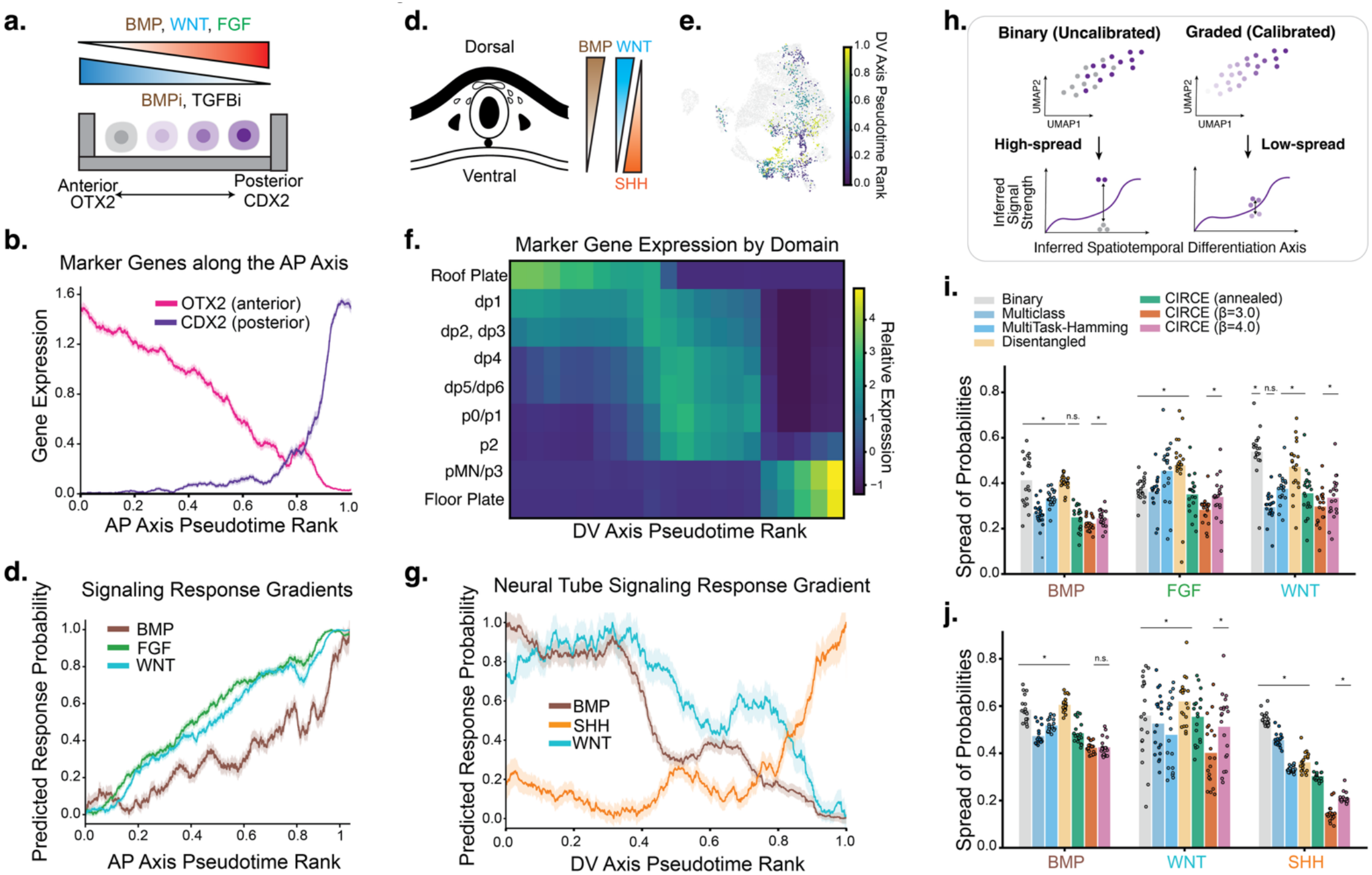
Graded signaling scores recover engineered and endogenous signaling gradients. **a**, Antiparallel signaling inputs along the anterior-posterior (AP) axis of a human endoderm differentiation model, imposed by a microfluidic system ^36^. **b**, Diffusion-pseudotime ordering of cells along the AP axis, validated by anterior *OTX2* and posterior *CDX2* expression. **c**, Predicted signaling response probabilities along the inferred AP axis. The probabilities for each pathway were independently min-max normalized to compare spatial shape. Spatial gradients were calculated by averaging cells within defined spatial bins along the pseudotime axis. **D**, Schematic for the spatial distribution of major signaling pathways in the developing neural tube. **E**, Inferred dorsal-ventral (DV) axis based on expression programs of canonical marker genes, using the mouse gastrulation atlas ^42^. **F**, Expression of marker gene programs across inferred cell-type domains in the developing neural tube, validating the spatial ranking by pseudotime. **G**, Predicted pathway activation scores along the DV axis reconstructed the known antiparallel gradients. **h**, Evaluation of model calibration based on the spread of inferred pathway activation scores. A well calibrated model produces predictions that have a tighter spread among cells in the same bin along the pseudotime axis. **i & j**, The spread of the inferred probability values for the relevant pathways in the microfluidic endoderm system (**i**) and the mouse neural tube (**j**). * indicates p < 0.05 by a two-tailed Wilcoxon ranked signed test with a Benjamini-Hochberg correction for false discovery.

CIRCE predictions successfully recovered the spatial gradients of signaling activities. We used Diffusion pseudotime to order cells along the anterior-posterior axis, which was marked by enriched *OTX2* expression anteriorly and *CDX2* expression posteriorly, confirming the expected orientation **(Fig. 4b)** (Haghverdi et al. 2016). CIRCE predictions varied continuously across this axis, with BMP, WNT, and FGF scores increasing toward the posterior in the expected direction, reconstructing the geometry of the imposed gradients from transcriptional state alone **(Fig. 4c, Supplementary Figs. 10-11)**. To avoid ambiguity in the ground-truth labels, we focused on signals that were activated by an externally provided agonist. The recovery of continuous, correctly-oriented gradients demonstrates that the model’s intermediate scores contain information about relative signaling activity strength, even though agonist concentration was absent from the binary treatment labels.

### The model identifies graded signaling organization *in vivo*

Finally, we tested whether the approach could recover signaling relationships in native tissues, where pathway activity was not experimentally imposed. The developing neural tube has characteristic morphogen gradients that define the dorsal-ventral axis: BMP and WNT specify the more dorsal cell types while an SHH gradient patterns the more ventral cell types **(Fig. 4d)** ^37–41^. We applied our models to scRNA-seq data from the developing mouse neural tube (E8.5) ^42^. We reconstructed the DV axis using cell fate markers for each of the differentiated cell types in the neural tube **(Table 2)**. We assigned dorsal and ventral scores based on the expression levels of the marker genes and constructed a pseudotime axis **(Fig. 4e-f, Supplementary Fig. 12)**. CIRCE recovered spatially graded WNT, BMP and SHH signaling activity profiles, with predicted WNT and BMP signaling activities receding as the SHH signaling activities increasing **(Fig. 4g, Supplementary Figs. 13-14)**.

**Table 2.** Neural tube cell type marker genes.

| **Cell type** | **Marker Genes** |
|---|---|
| **Roof Plate** | *LMX1A, LMX1B, MSX1, MSX2, WNT1, GDF7* |
| **dp1** | *ATOH1* |
| **dp2** | *NEUROG1* |
| **dp3** | *OLIG3, IRX3* |
| **dp4** | *GSX1, GSX2, ASCL1* |
| **dp5** | *GSX1, ASCL1, PAX7* |
| **dp6** | *DBX2, PAX7* |
| **p0** | *DBX1* |
| **p1** | *DBX2, NKX6-2* |
| **p2** | *NKX6-1, IRX3, FOXN4* |
| **pMN** | *OLIG2* |
| **p3** | *NKX2-2* |
| **Floor Plate** | *FOXA2, SHH, ARX* |

Notably, compared to other models, CIRCE appeared to be better calibrated. To evaluate calibration *in* vivo, we quantified the spread of predicted probability values within each averaging bin **(Fig. 4h)**. A model with a lower spread of predicted probability values is more reliable at recapitulating known biological signaling strength at the single-cell level. Notably, when compared to other models, CIRCE predictions were tighter around all points along both the microfluidic-imposed AP axis and neural tube DV axis **(Fig. 4i, j)**.

We also evaluated embryonic somitic mesoderm, in which signaling activities also vary along the body axis^42^. Relative to models trained with hard labels, CIRCE produced spatially extended, continuous signaling profiles across the expected cell-state progression **(Supplementary Fig. 15a)**. Like the neural tube case, CIRCE models produced much tighter distributions at each point along the inferred AP axis **(Supplementary Fig. 15b)**. These results demonstrate that models calibrated with the combinatorial structure in the training data can reveal a more graded signaling organization than models without calibration, even in biological contexts that differ substantially from the perturbation screen used for training.

## Discussion

Binary combinatorial screens do not merely produce binary responses to each perturbation; instead, they contain a hidden response space that varies continuously in magnitude across all cells. We show that this structure can be used to learn graded representations of cellular responses to perturbations without requiring continuous experimental labels. Crucially, in a combinatorial screen, each perturbation is observed across multiple cellular contexts, generating a range of response magnitude that has inherently encoded structure. We exploit this structure by using Boltzmann smoothing over Hamming distances to encourage predicted probability distributions to be locally coherent in combinatorial space. Consequently, predictions that differ from the observed condition by a single pathway receive more credit than those separated by multiple pathways. Marginalizing the resulting joint distribution produces pathway scores that are less polarized, better calibrated, and aligned with variations in signaling levels across tissues. Thus, combinations become not a classification burden, but a source of calibration.

Our results further suggest that uncertainty can capture meaningful biological signals. While the model is trained on discrete identity labels, cellular responses themselves do not necessarily segregate into hard categories. Our active-learning and gene-response analyses indicate that much of the resulting uncertainty reflects irreducible ambiguity due to a continuum of response states rather than inadequate model training. Therefore, we interpret the prediction scores as relative measures of pathway activity. Future datasets containing matched ligand doses, direct pathway reporters, or phosphoproteomic measurements will be needed to map these scores onto absolute activity levels.

This structured-label smoothing approach also has practical implications for experimental design. Our finding suggest that models may scale more effectively by training on diverse combinations than by collecting ever-larger numbers of cells from a narrow set of conditions or exhaustively titrating each input. Combinatorial perturbation screens could therefore be designed not only to identify interactions, but also to span the response space required for calibration. An important direction for future work will be to determine which combinations are most informative for calibration and out-of-distribution prediction.

More generally, we show that distinct training objectives can leverage seemingly unrelated properties of biological data. Even with a simplified representation of perturbation relationships, such as by using the Hamming distance between combinations, our models learned both signal identity and graded response states. A useful next step would be to define or learn pathway-specific distance measures from perturbation data or prior network knowledge, allowing the label geometry to capture synergistic, antagonistic, and directional interactions. Additionally, incorporating key inhibitory molecules, such as signaling pathway antagonists, into a probabilistic framework should further improve the ability to fine-tune predicted signaling histories or spatial distributions.

In summary, our structured-label framework repurposes binary combinatorial perturbations to learn continuous representations of cellular states. By recovering graded pathway activities without dose-resolved training annotation, it provides a general route towards modeling dose-dependent cell-state transitions. Applied to developmental signaling contexts, these models could guide the differentiation of stem cells into desired cell types, and the design of organoids with increasingly precise multicellular organization. More broadly, extending this approach to small-molecule drug perturbation screens could enable predictions of how drug doses shape target-pathway activity, supporting the rational design of dose-optimized therapies.

## Acknowledgements

The authors would like to thank Miram Meziane and Mark Greenwood for helpful discussions. NTH would additionally like to thank the generous support of the National Science Foundation Graduate Research Fellowship Program. P.L. discloses support for the research of this work from National Institute of Health (DP2HD108777, R00HD087532), Allen Family Philanthropies, Eugene Bell Career Development Professorship, and Centurion Foundation.

## Author contributions

N.H. conceived and executed the research. P.L. and N.H. supervised the research. S.M. and C.L. performed key experiments and tests..

## Methods

### Perturbation screen data processing

For the combinatorial signal perturbation screens ^13,25^, single-cell RNA-sequencing data were processed using scanpy. Raw counts were library-size normalized to 10000 counts per cell and log-transformed. Highly variable genes were selected using standard dispersion-based criteria (min_mean = 0.0125, max_mean = 3, min_dispersion = 0.5). Cells were drawn from six experimental batches from two combinatorial signaling ligand screens, both of which include six key developmental signaling pathways (BMP, FGF, RA, SHH, TGFB, WNT).

### Hamming distance and Transcriptional Distance Comparison

To compute the relationship between Hamming distance and transcriptional distance between cells, pairs of cells were chosen randomly from the dataset with ground truth signaling labels. The Hamming distance between the binary strings of signaling combinations was computed between the cells. The transcriptional distance was computed through three different metrics in the highly-variable gene space: cosine, Euclidean and Wasserstein.

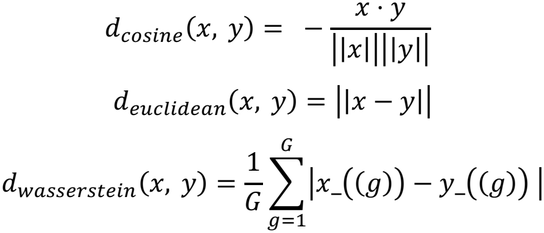

Where G is the number of variable genes (G=2000).

For the neighborhood-based analysis, the k-nearest neighbor graph was constructed for varying values of k and the average Hamming distance between cell signaling combinations was calculated.

### Variational autoencoder backbone

All classifiers were built on a shared VAE backbone using scVI as implemented in scvi-tools ^19,28^. The encoder maps gene expression counts to a Z-dimensional latent space under a negative binomial likelihood and batch corrected latent representations are obtained by conditioning the decoder on one-hot batch covariates. The training objective is the evidence lower bound (ELBO).

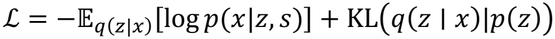

Here, s is the batch covariate. Models were trained for 40 epochs with the Adam optimizer (η = 5 × 10; ^−4^, *γ* = 1 × 10; ^−6^, batch size of 64, and a linear KL warm-up over 3000 gradient steps). Gradient norms were clipped at 1.0 to prevent instability.

### Signaling state classification benchmarking

Pathway-state classifiers were implemented by fine-tuning scANVI models on the scVI backbone. Classifiers were fine-tuned for five epochs using early stopping (patience = 10), a batch size of 256, and gradient clipping at 1.0. We compared four baseline formulations with CIRCE. (1) The binary formulation used six independent pathway classifiers. Pathway-specific architectures were selected by cross-validation over depth L in {1,2, hidden width H in {512,2048, and latent dimension Z in {16,32,64. (2) The conventional multiclass formulation used a single 64-class softmax classifier (L = 2, H = 1,024, Z = 64). (3) The multitask formulation used a shared latent representation with six binary cross-entropy heads. (4) The joint formulation optimized both multiclass and binary objectives. CIRCE used the same architecture as the conventional multiclass model but was trained with the Hamming-Boltzmann soft-label objective described below.

For multiclass models, the score for pathway m was calculated by summing the predicted probabilities of all combination classes in which pathway m was active. This marginalization yielded one score between zero and one for each pathway and cell.

### Hamming-Boltzmann soft-label loss

Each of the 64 co-stimulation conditions was represented as a six-bit vector. For a cell with observed class y, the soft target assigned to class *c*_*j*_ was

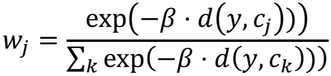

Where *d*_*H*_ is the Hamming distance between two signal combination strings and *β* is an inverse-temperature parameter. The classification objective was the cross-entropy between this soft target and the predicted class distribution:

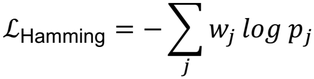

During training, beta was annealed exponentially from 3.0 to 10.0 or held constant at values 3.0 or 4.0. This annealing schedule provided broader smoothing early in optimization and increasingly concentrated the target distribution around the observed class later in training.

Cross entropy with label smoothing was performed by subtracting either a fixed *ϵ* from the target labels, an *ϵ* divided by the hamming distance of the nearest combination, or an *ϵ* divided by the entropy computed from the distribution of outputs of an MC dropout estimate. *ϵ* was set to {0.05, 0.1, 0.2}.

### Uncertainty estimation

Predictive uncertainty was estimated using Monte Carlo dropout (MC) and ensembles of independently initialized models ^31^. For Monte Carlo dropout, dropout layers were kept activate at inference time and each cell was passed through the trained classifier 10 times (T), yielding class distributions over the 64 signaling combinations {*p*_1_, *p*_2_, . . . *p*_*T*_}. Predictive entropy was calculated from the mean class distribution, 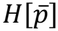. We define 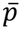 as 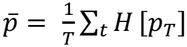 . This was decomposed into aleatoric uncertainty, the mean entropy of the individual passes 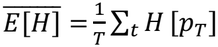, and mutual information between predictions and model parameters, *I*= *H*[*p*] − *E*[*H*].

For the binary model, the entropies for each of the six pathways were summed, which equals the joint entropy over the 64 combinations under independence and places all models on a common scale. For the ensemble estimator, the same decomposition was applied across the five models (N) in place of dropout passes {*p*_1_, *p*_2_, . . . *p*_*N*_}. This vector of probability values can also be used to compute the mean entropy, mutual information and aleatoric uncertainty in the same way. All uncertainties were computed on held-out test cells from the raw probability values.

### Active Learning

Bayesian Active Learning by Disagreement (BALD) was implemented by using the Monte Carlo dropouts and ensembles to calculate the difference between the entropy of the mean prediction and the mean entropy of individual stochastic predictions ^43^. To test whether uncertainty identified informative missing examples, one experimental batch was held out in each cross-validation fold. 20% of held-out cells were reserved for testing, and the remaining pool of 80% were successively acquired during the experiments. The classifier was initialized with acquisition-pool labels masked. In each of four rounds, 15% of the remaining pool was selected, labels were revealed, and the classifier was retrained.

We compared two acquisition strategies: BALD and random sampling. In random sampling, cells were chosen randomly from the remaining pool. Random sampling was repeated with three seeds and followed the same acquisition ratios as BALD. The same analysis was performed for six held-out-batch folds and for an in-sample random split.

### Synthetic Data Generation

Counts were simulated for all 64 possible combinations of the six pathways from a negative-binomial model in which each pathway increased the log-mean expression of its own 40-gene marker block alongside 200 non-responsive genes. Aleatoric uncertainty was introduced by flipping the recorded labels with a probability of 0.3, independent of expression. Epistemic uncertainty was simulated by adding additional cells which expressed a 120-gene program that was not present in any of the other cells. All models were trained on the labeled cells with the same architectures and settings as the other analysis. 30% of labeled cells were held out for testing. Uncertainty was decomposed with 15 MC-dropout passes into aleatoric and epistemic components.

### Evaluation Metrics

Categorical performance was evaluated using exact-match accuracy and normalized Hamming distance. Exact-match accuracy was the fraction of cells for which all six pathway labels were predicted correctly. Normalized Hamming distance was the fraction of pathway-level mismatches per cell. Pathway-level performance was additionally assessed using accuracy, area under the precision-recall curve, area under the receiver operating characteristic curve, and F1 score.

Calibration was quantified using expected calibration error (ECE) by grouping predictions according to confidence and calculating the weighted difference between confidence and empirical accuracy across bins. For multiclass models, calibration was evaluated both for the complete combination and for marginalized pathway scores. Intermediate pathway scores were defined as values between 0.1 and 0.9; this threshold was used descriptively to compare the polarization of model outputs and was not treated as a biochemical activity range.

### Monotonicity of Response Gene Signaling Association

To compute the monotonicity of a set of genes, cells were grouped into deciles of stimulation probability for each signal, averaged over five training seeds and the mean log-normalized expression of the gene set was taken for each decile. The monotonicity was computed as the Spearman Correlation between gene expression and decile index. To compute empirical p-values, a null distribution was computed as sets of 200 randomly selected gene sets of equal size drawn from the 2000 HVGs. The Spearman correlation coefficient is calculated as follows:

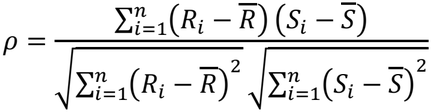

This is where *R*_*i*_ is the decile of the stimulation score and *S*_*i*_ is the percentile of the gene expression score.

### Analysis of graded signaling systems

For the microfluidic endoderm dataset, diffusion pseudotime was used to order cells along the inferred anterior-posterior axis. Diffusion pseudotime generates a diffusion map from the nearest-neighbor graph (k = 15) transition matrix and each cell’s pseudotime is its diffusion distance from a designated root cell ^44^. Diffusion pseudotime was computed using scanpy.tl.diffmap in scanpy 1.10.0. These trajectories were rooted based on marker genes relevant for each cell type. Axis orientation was verified using *OTX2* and *CDX2* expression. Pathway scores were predicted for individual cells and summarized within each bin along the pseudotime axis. Where indicated for comparison of spatial shapes, each pathway profile was min-max normalized independently. For neural tube differentiation, BMP, WNT and SHH scores were ordered along the dorsal-ventral axis with classic differentiation markers (Table 2). The same trained framework was applied to embryonic somitic mesoderm, and predicted activities were visualized across the inferred developmental axis.

### Tightness of a Gradient Prediction

To assess the consistency of predictions across a spatiotemporal gradient, cells were ordered along an axis computed with pseudotime and validated by marker gene expression. They were then divided into 20 equal-count positions. At each position, k, the tightness was computed as the sample standard deviation normalized by the Bhatia-Davis inequality ^45^:

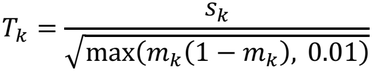

## Data availability

All data used in this study are previously published and publicly available. These datasets include a combinatorial signal screen on mouse embryonic stem cell (GSE289836), three combinatorial signal screens on human embryonic stem cells (GSE327767), a mouse embryo gastrulation atlas (GSE 87038), and microfluidic patterned human foregut model (GSE285075).

## Code availability

All code used in this study is available at: https://github.com/nhutchins627/circe-signaling.

## Supplementary Figures

**Supplementary Figure 1.**
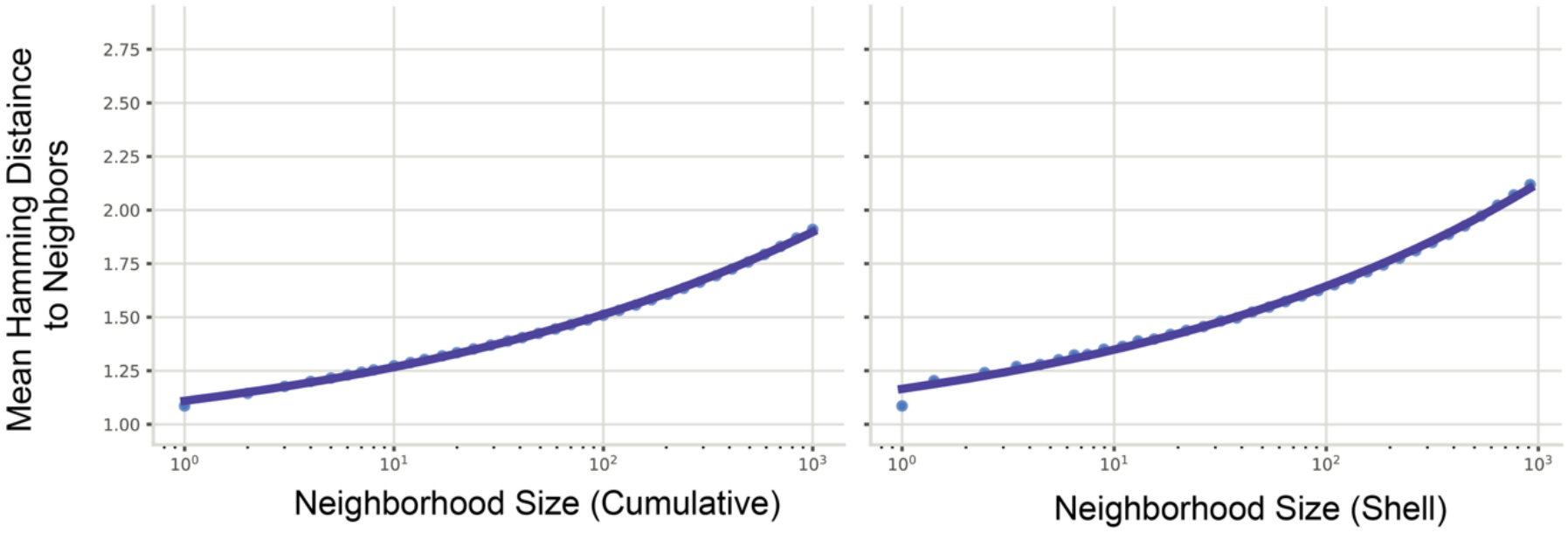
More distant neighbors have larger Hamming distance between their signaling combinations. We survey different choices of the number of nearest neighbors to consider, k. Left, the mean hamming distance between neighbors for a given neighborhood size. Right, the mean hamming distance between neighbors for a given neighborhood size k, excluding all those cells that are present in neighborhoods of size k-1.

**Supplemental Figure 2.**
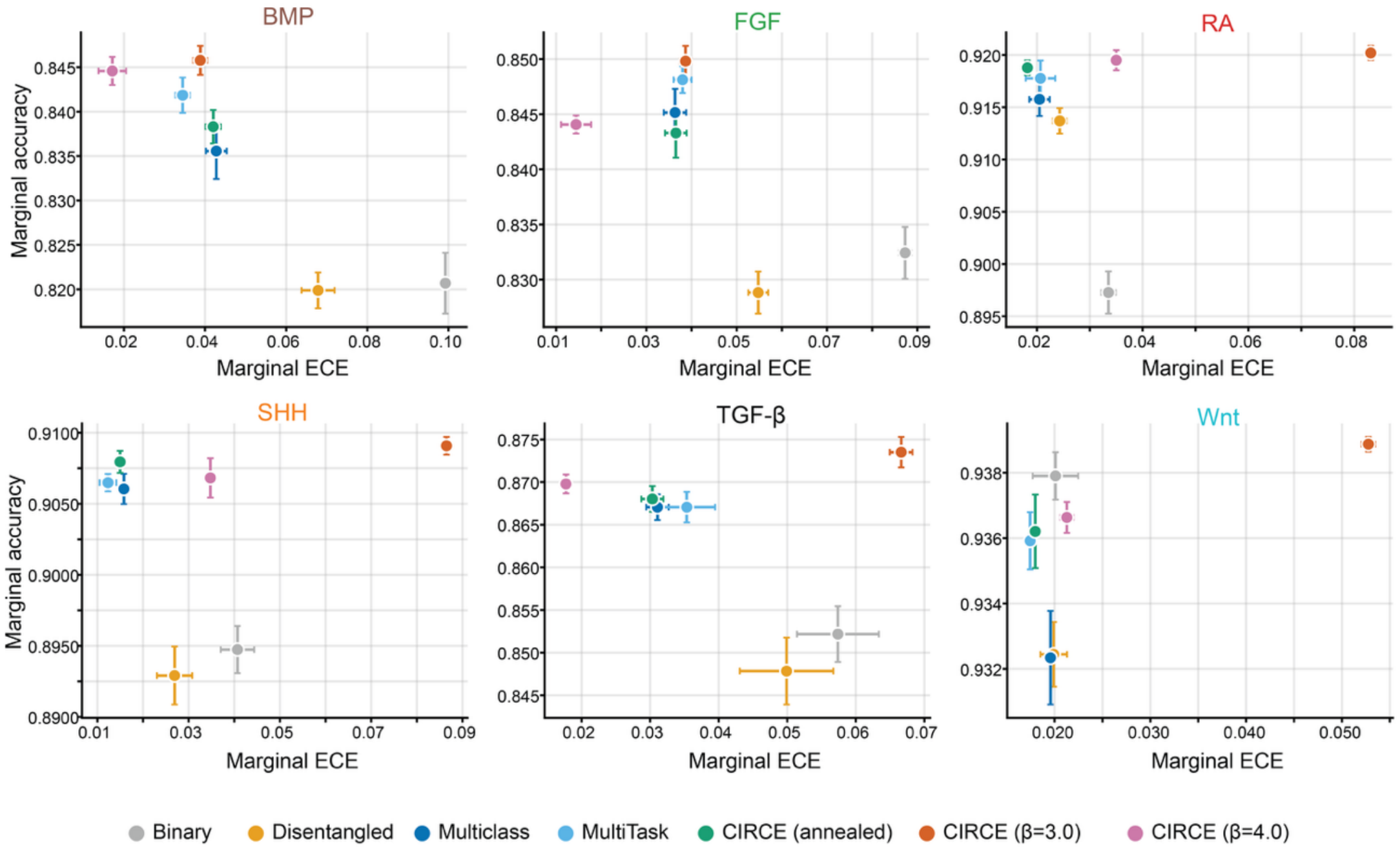
ECE and accuracy comparison among models across all pathways. The marginal accuracy is the per-signaling-pathway accuracy of a combinatorial prediction. Error bars denote standard deviations of ECE and accuracy. Models that fall in the upper-left quadrant have high accuracy and optimal ECE. In general, CIRCE ranks highly amongst the models across all pathways.

**Supplemental Figure 3.**
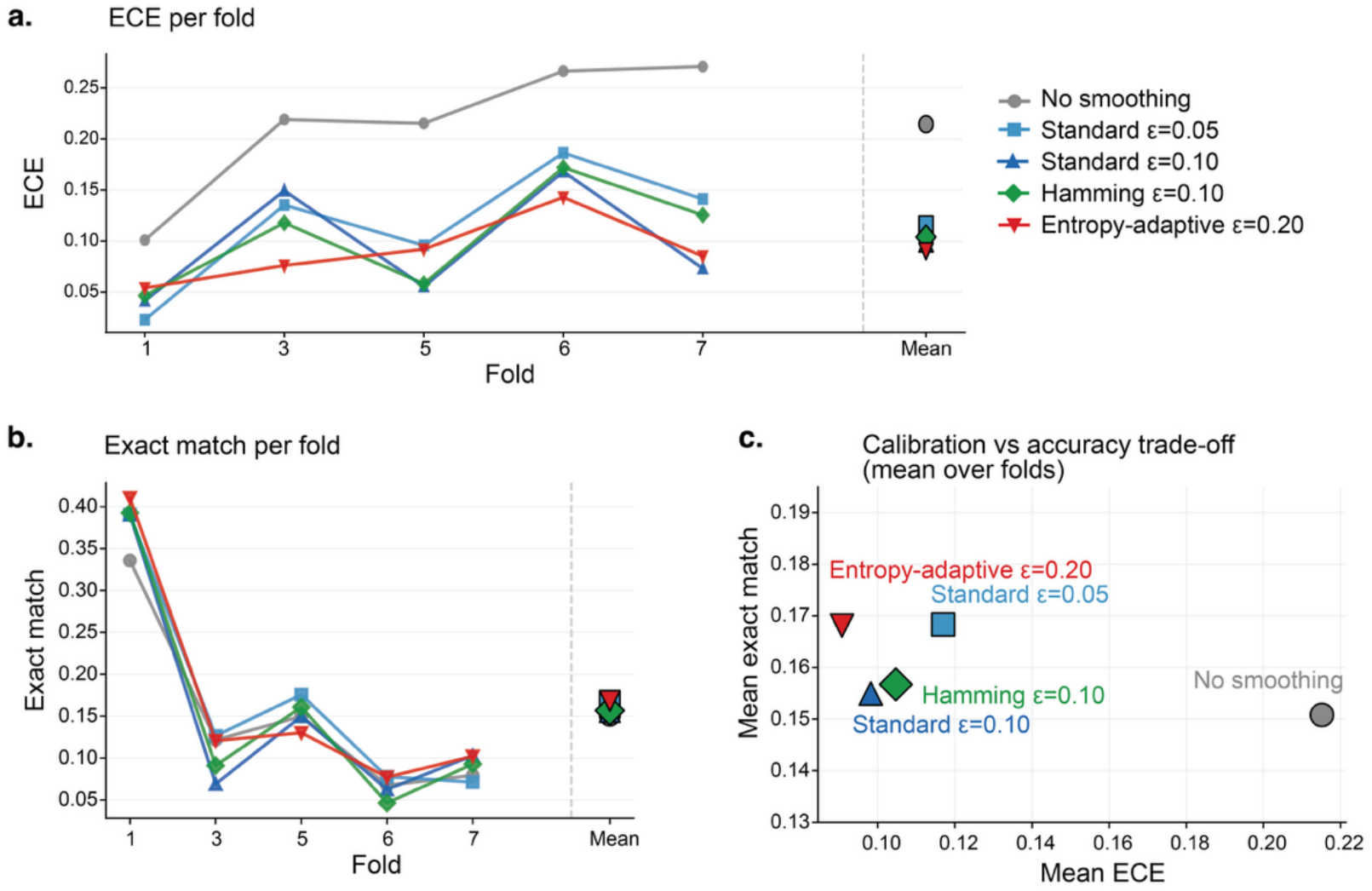
Label smoothing improves calibration without reducing exact-match accuracy. **a**, ECE of Binary model with generic label-smoothing methods applied, including standard, Hamming-based, and entropy-adaptive label-smoothing. ε represents smoothing strength. Folds represent active learning acquisitions of unseen training data. In standard smoothing, cells have their target labels smoothed by an ε value of 0.05 or 0.1. In the Hamming-based smoothing, cells with further Hamming distance relative to seen combinations from the reference cells have their labels smoothed by an ε value of 0.1. In entropy adaptive, labels are smoothed by 0.2 divided by the predictive entropy of that cell calculated via MC dropouts. **b**, Assessing the effectiveness of label smoothing with the exact-match accuracy across folds. **c**, Label smoothing generally improves model accuracy and ECE, compared to no smoothing. Mean exact match and mean ECE are the average of the active learning folds.

**Supplemental Figure 4.**
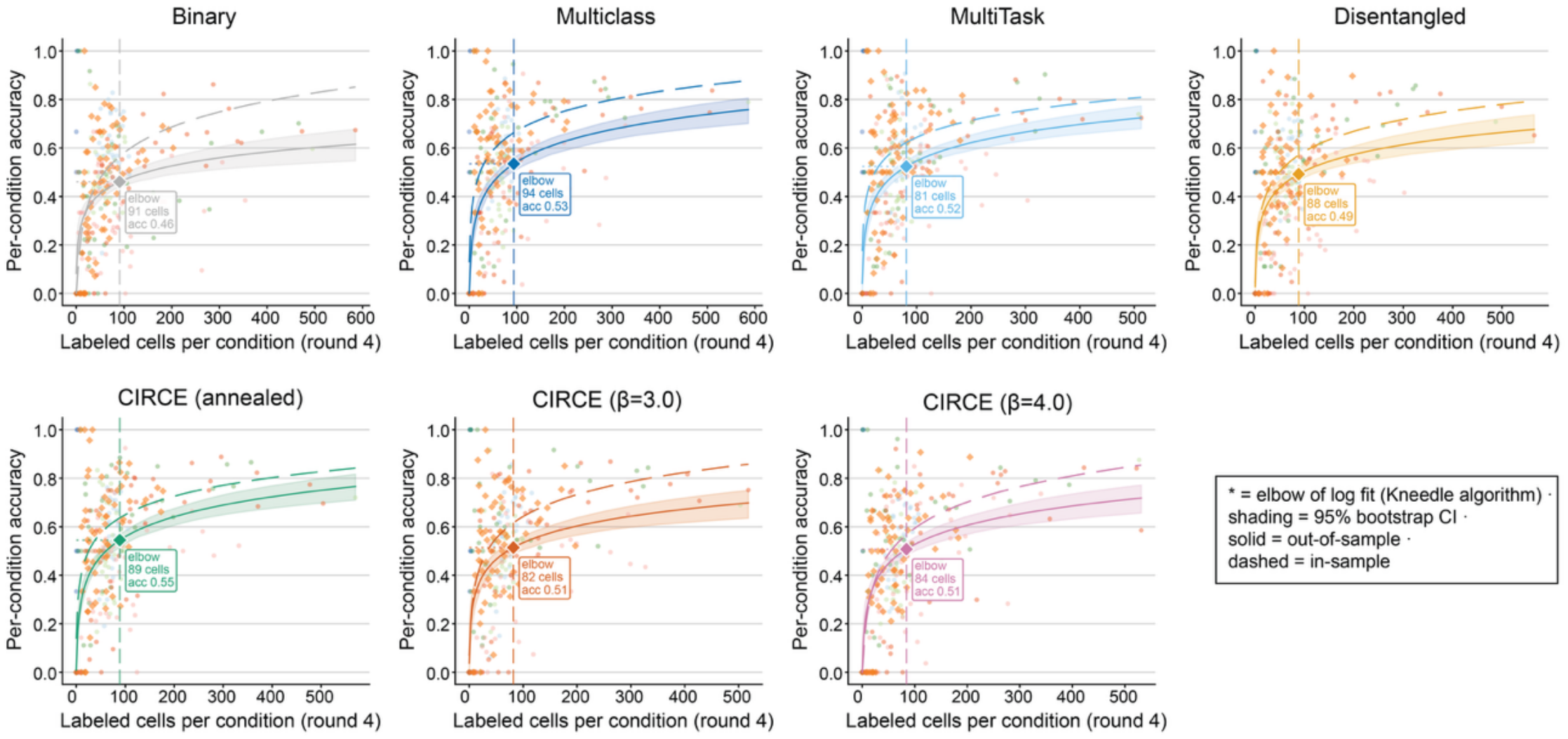
Performance saturates with increasing cells per combination. Model accuracy for each combination increases with the number of labeled cells for each perturbation. For each model, the exact match accuracy consistently begins to plateau in the <100 cell regime, suggesting that models converge to their maximum combination-level accuracy with a relatively small sample size. The data was partitioned into distinct out-of-sample (OOS) folds based on cell type. In each case, the OOS represents the average across held-out folds when evaluated on unseen cell types and in-sample represents the average across random splits of the data. The elbow was computed with the Kneedle Algorithm as the point of maximum curvature.

**Supplemental Figure 5.**
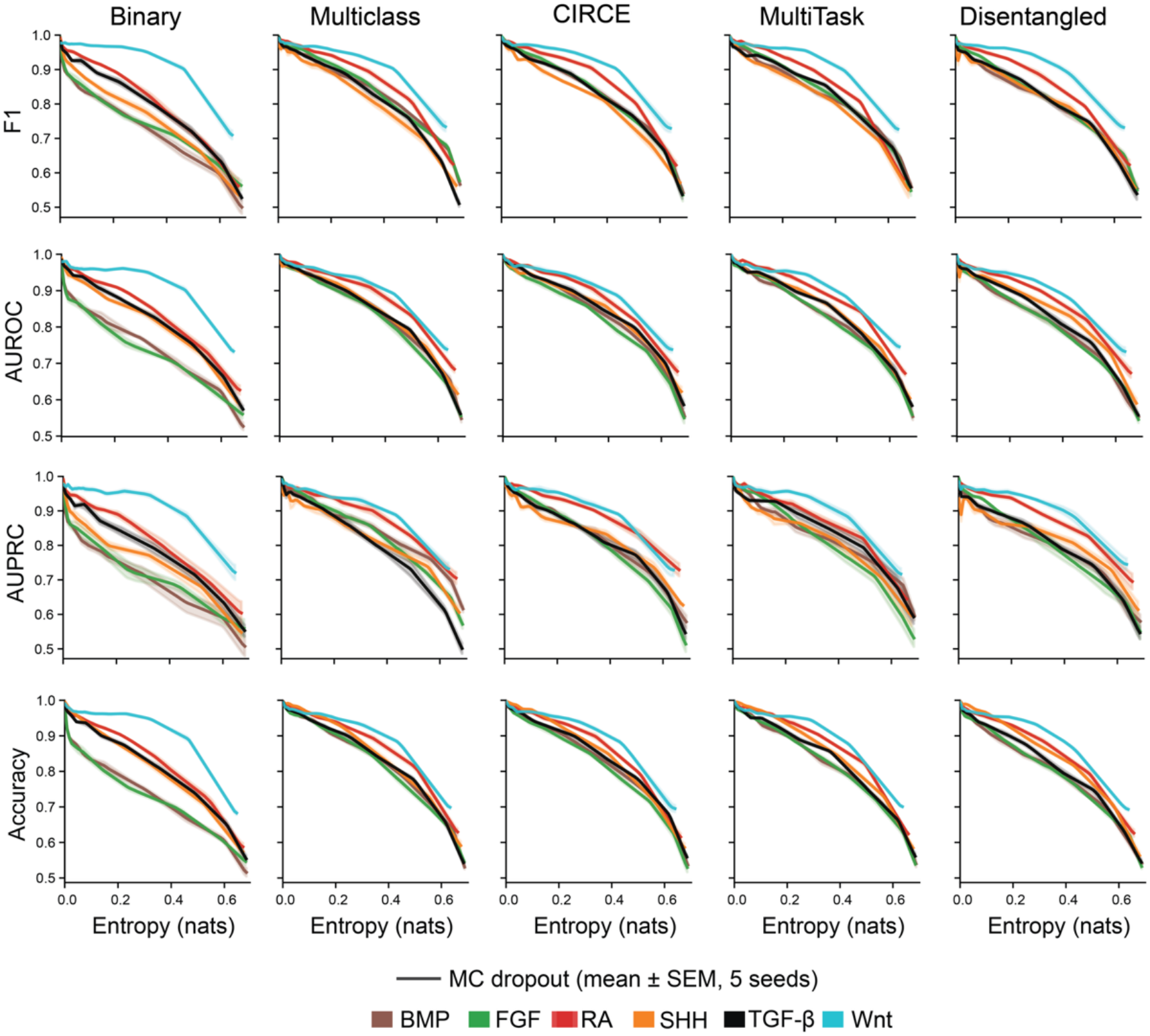
Predictive uncertainty is associated with reduced classification performance. Each model is shown with its predictive uncertainty by MC dropouts. Across all models, accuracy, AUPRC, AUROC, and F1 score decrease with increasing predictive entropy across evaluated models. This validates the metric as informative for model performance. Accuracy was computed as the number of matching class predictions.

**Supplemental Figure 6.**
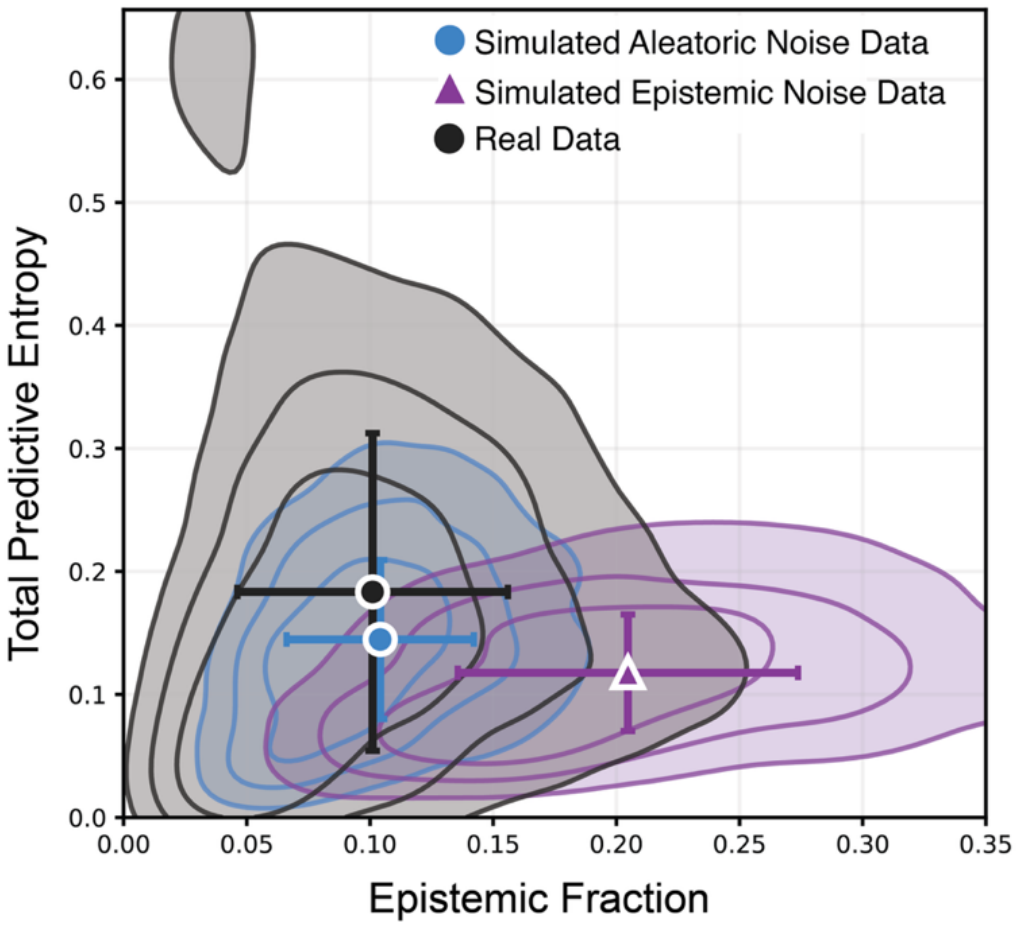
Uncertainty decomposition distinguishes aleatoric- and epistemic-dominated uncertainty. Total predictive entropy (y-axis) and BALD-decomposed epistemic fraction (x-axis) for synthetic and perturbation datasets in arbitrary units. The decomposition of the noise in models trained on the real screening data (*black*), an aleatoric simulated case (*blue*), and an epistemic dominant case (*purple*). The closer alignment between the simulated aleatoric dominated-case and real-data decomposition indicates that aleatoric noise is dominating in our experiments.

**Supplemental Figure 7.**
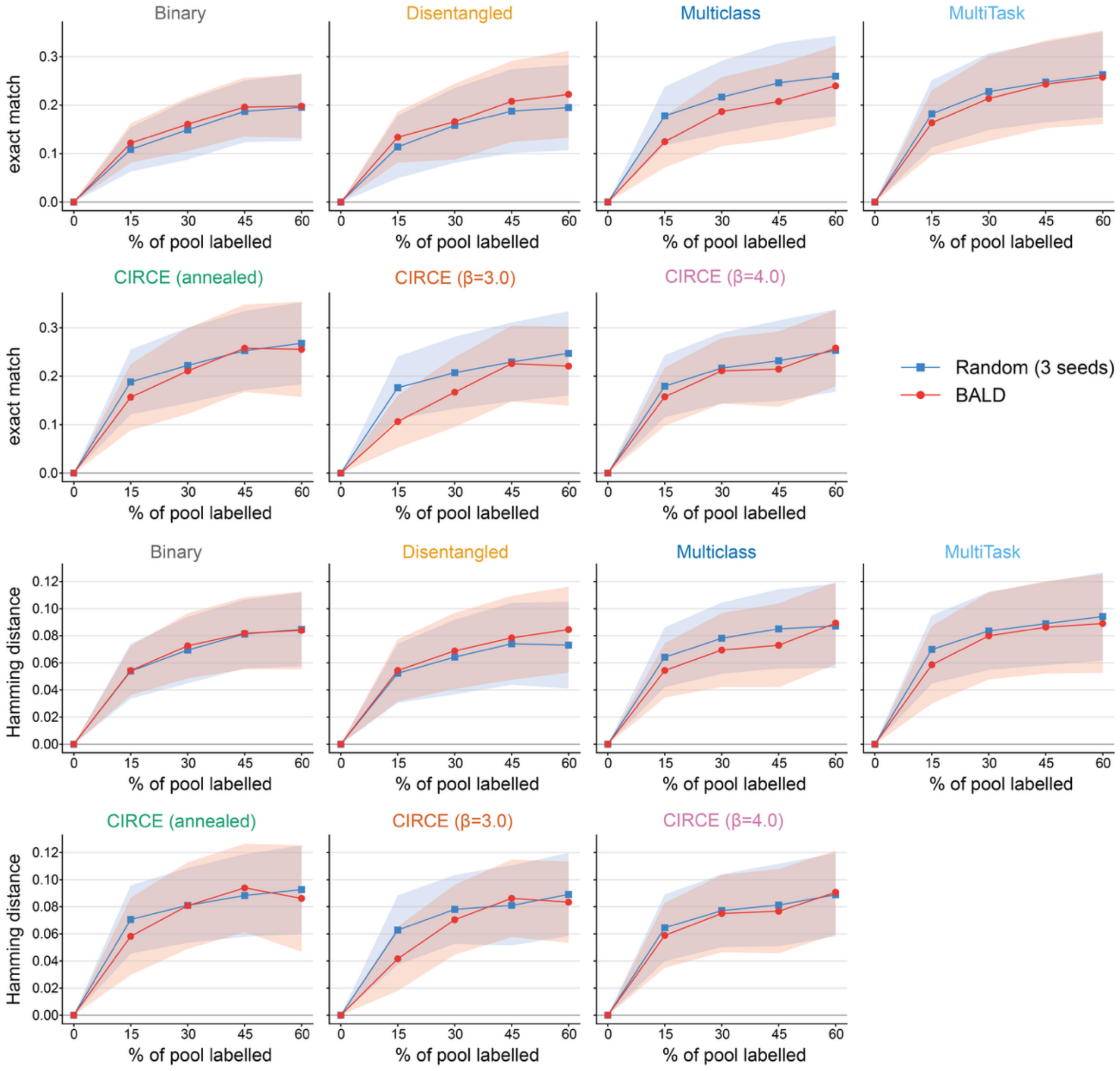
Exact match and Hamming distance across active-learning acquisitions. Each active learning fold is shown as a point on the x-axis and the y-axis is the performance metric of interest, either the Hamming distance from the true label or the exact match accuracy. Hamming distance and exact match across acquisition rounds were normalized across data splits and cell types. No significant difference was observed between the BALD (*red*) and random sampling (*blue*) (Wilcoxin rank-signed test with multiple hypothesis correction, p > 0.05, n.s.). Solid lines denote the average across data splits, and shaded areas denote variance.

**Supplemental Figure 8.**
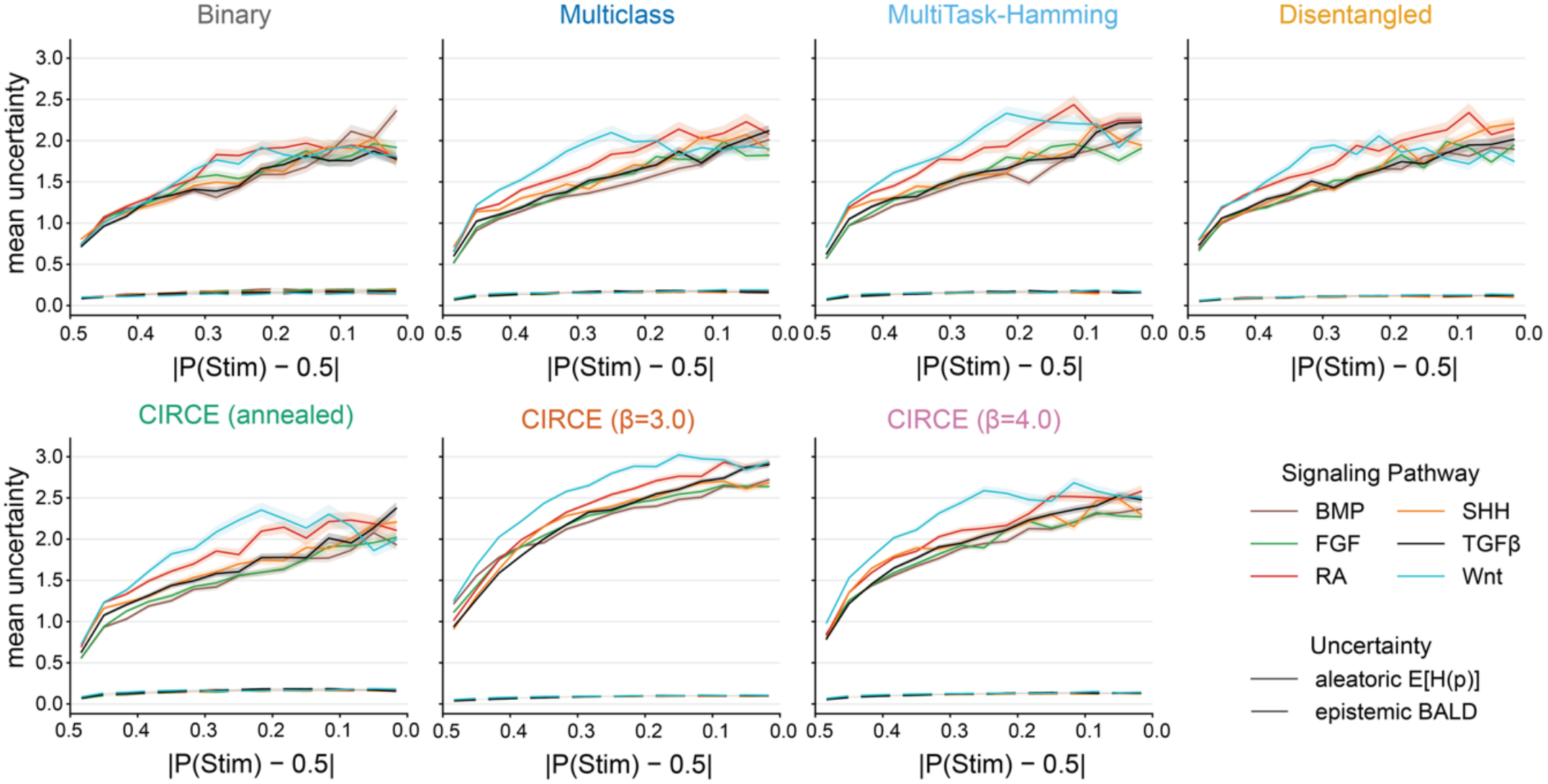
Uncertainty is highest for intermediate pathway scores. Estimated uncertainty as a function of predicted pathway activation scores for each signaling pathway across all models. Uncertainty was further decomposed into epistemic uncertainty (*dashed line*) and aleatoric uncertainty (*solid line*), displayed as average across folds (*lines*) with SEM (*shaded areas*). X axis represents the absolute deviation of predicted pathway activation scores from the intermediate probability value 0.5, with the highest deviation at the origin (predicted activation score of either 0 or 1).

**Supplemental Figure 9.**
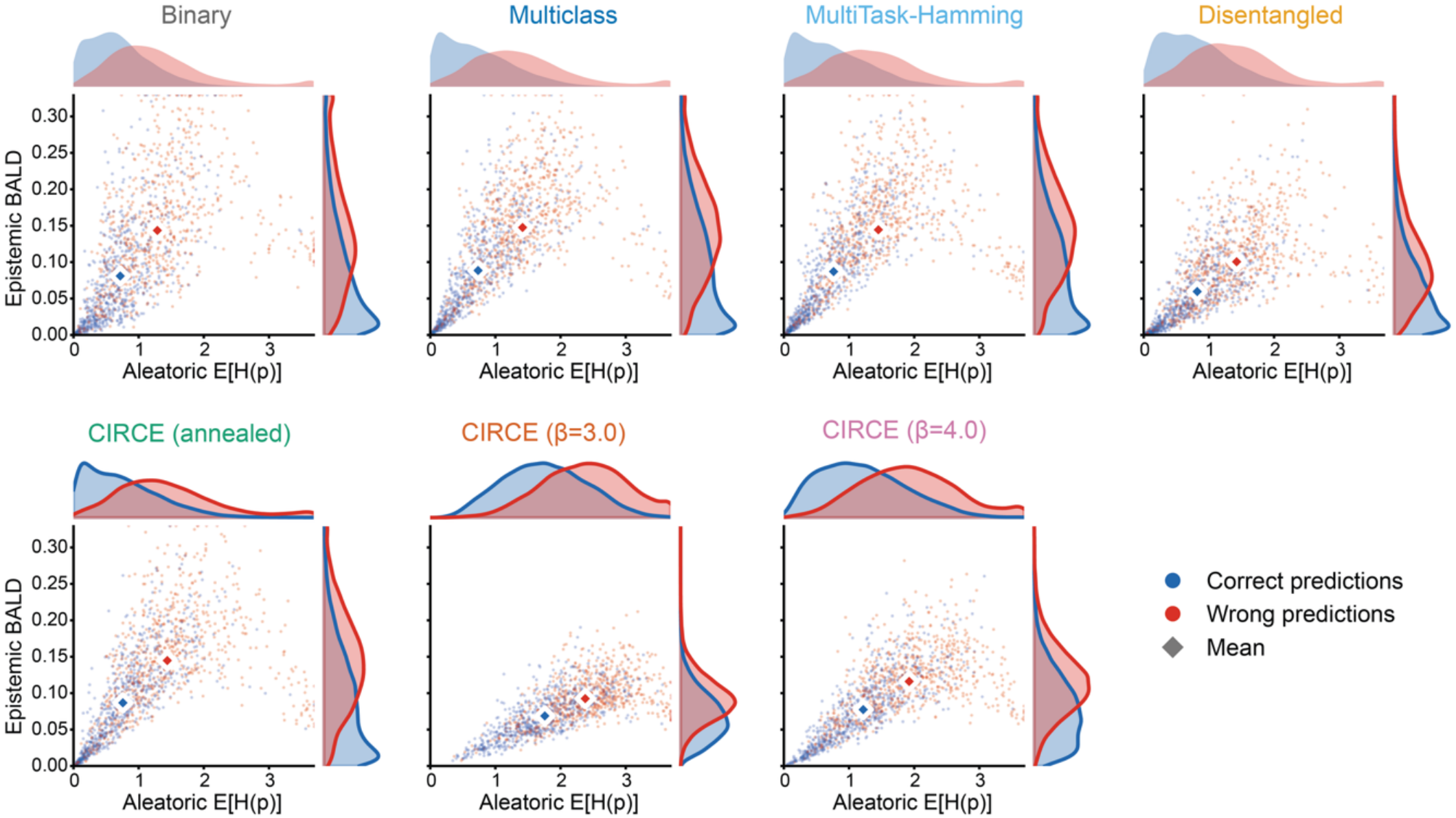
Uncertainty decomposition across model classes. Estimated aleatoric and epistemic components for correct and incorrect predictions, classified based on their exact match accuracy. Across all models, correct predictions have lower aleatoric and epistemic uncertainty according to BALD, relative to incorrect predictions .

**Supplemental Figure 10.**
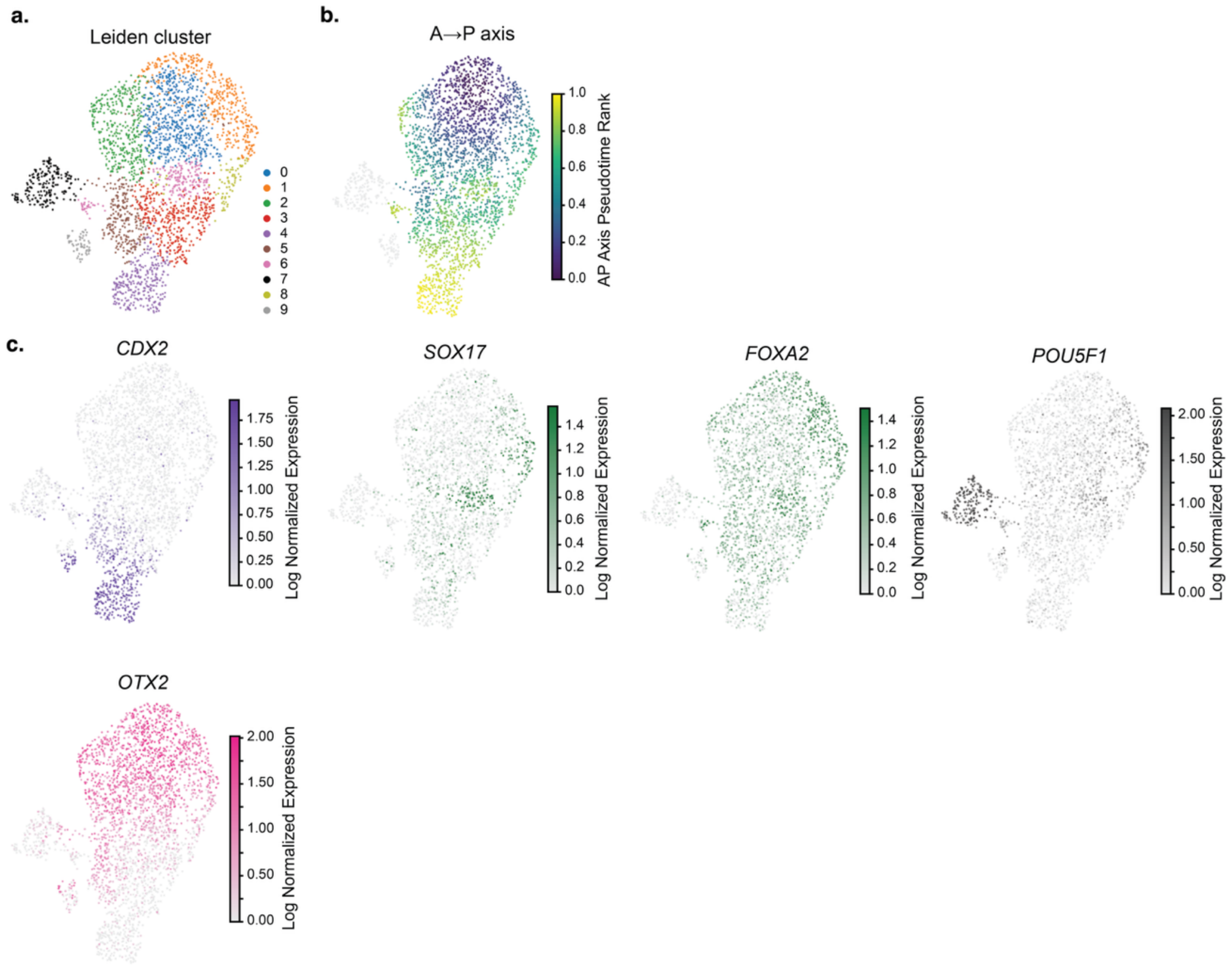
Marker gene expression in the microfluidic foregut model. **a**. Inferred leiden clusters through standard nearest neighbor-graph based clustering. **b**. Anterior-posterior scores is plotted as a function of gene sets marking spatial position. **c**. Expression of canonical cell type markers with known anterior-posterior patterns.

**Supplemental Figure 11.**
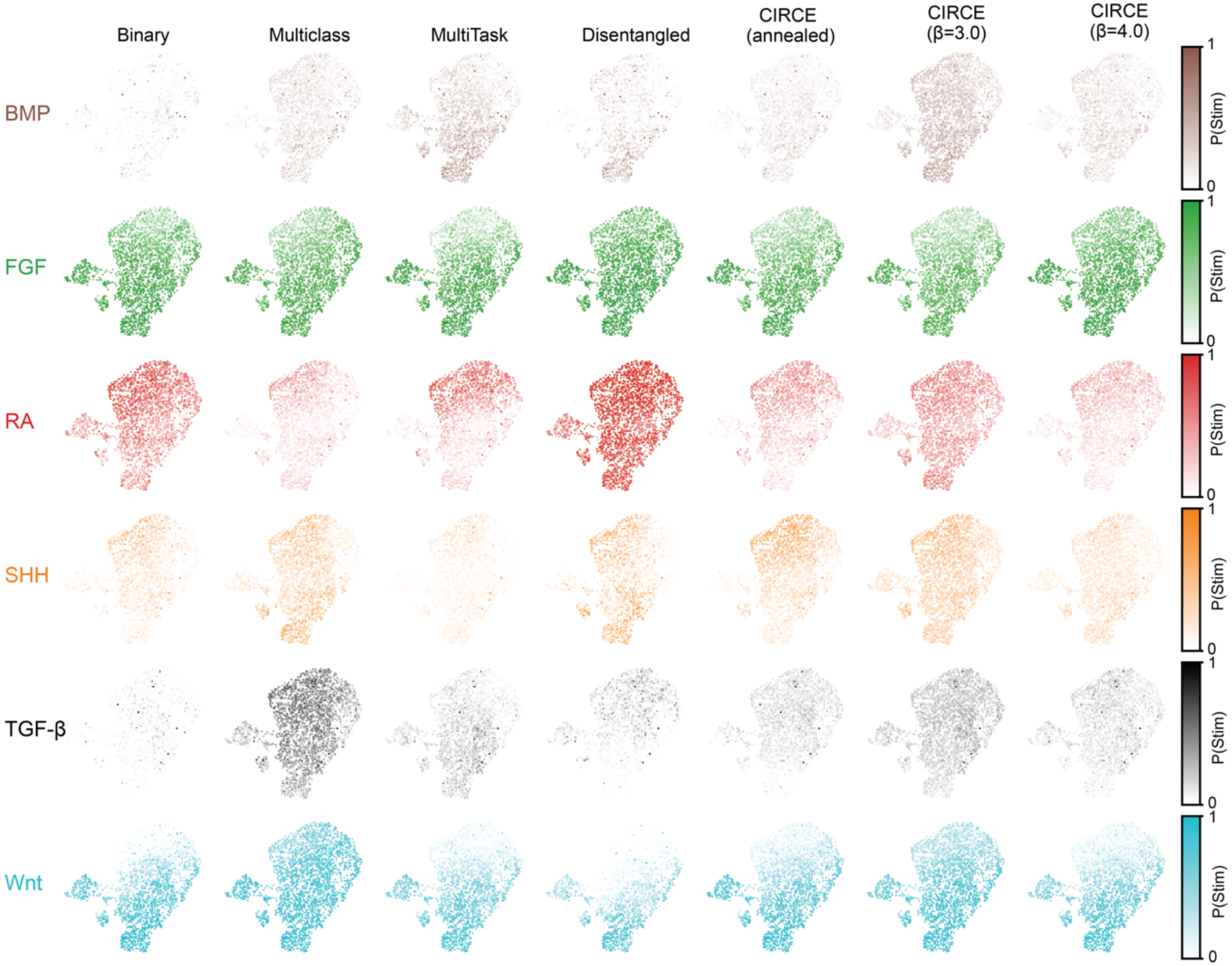
Predicted signaling pathway activation scores in the microfluidic foregut model. Cross model comparison of pathway activation plotted on UMAP from Supp. Fig. 10. Opacity of color corresponds to strength of signaling pathway activation for each of the individual signals.

**Supplemental Figure 12.**
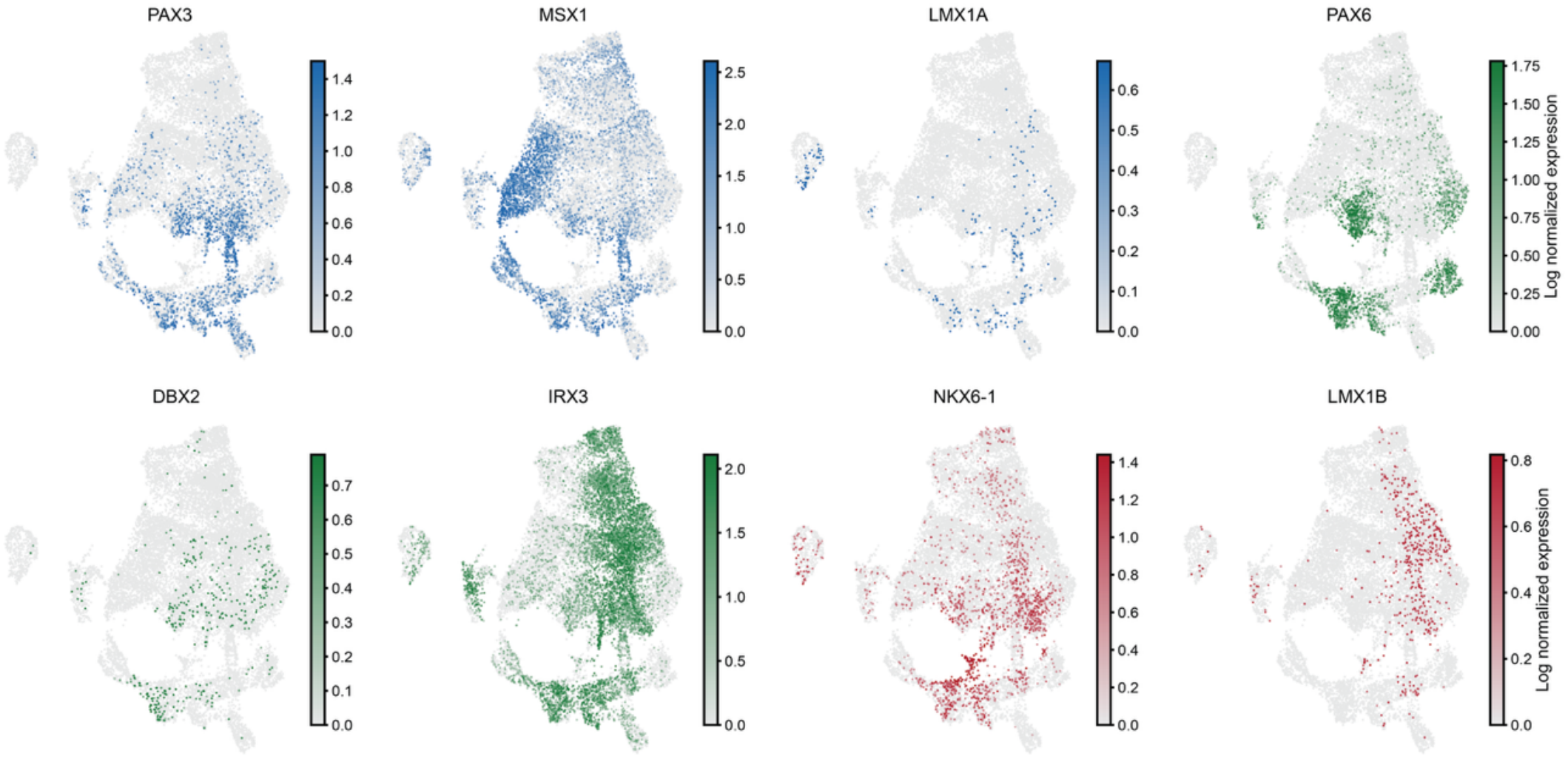
Marker gene expression in the developing neural tube. Canonical cell type marker genes for the dorsal-ventral axis in the early neural tube *(Dorsal: PAX3, MSX1; Roof Plate: LMX1A; Intermediate: PAX6, DBX2, IRX3; Ventral: NKX6-1; Floor: LMX1B)*. UMAPs were computed with standard nearest neighbor processing.

**Supplemental Figure 13.**
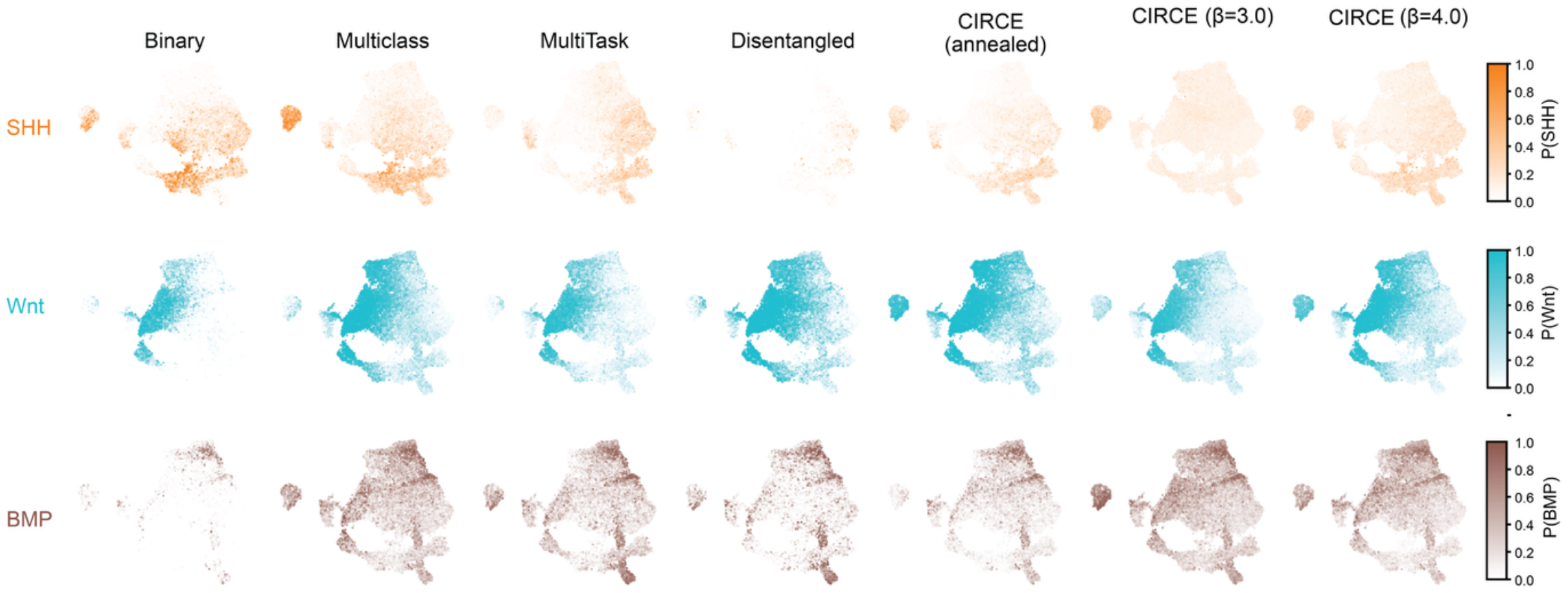
Predicted signaling pathway activation scores in the developing neural tube. Cross model comparison of pathway activation on the same UMAPs as in Supp. Fig. 12. Opacity of color corresponds to strength of signaling pathway activation.

**Supplemental Figure 14.**
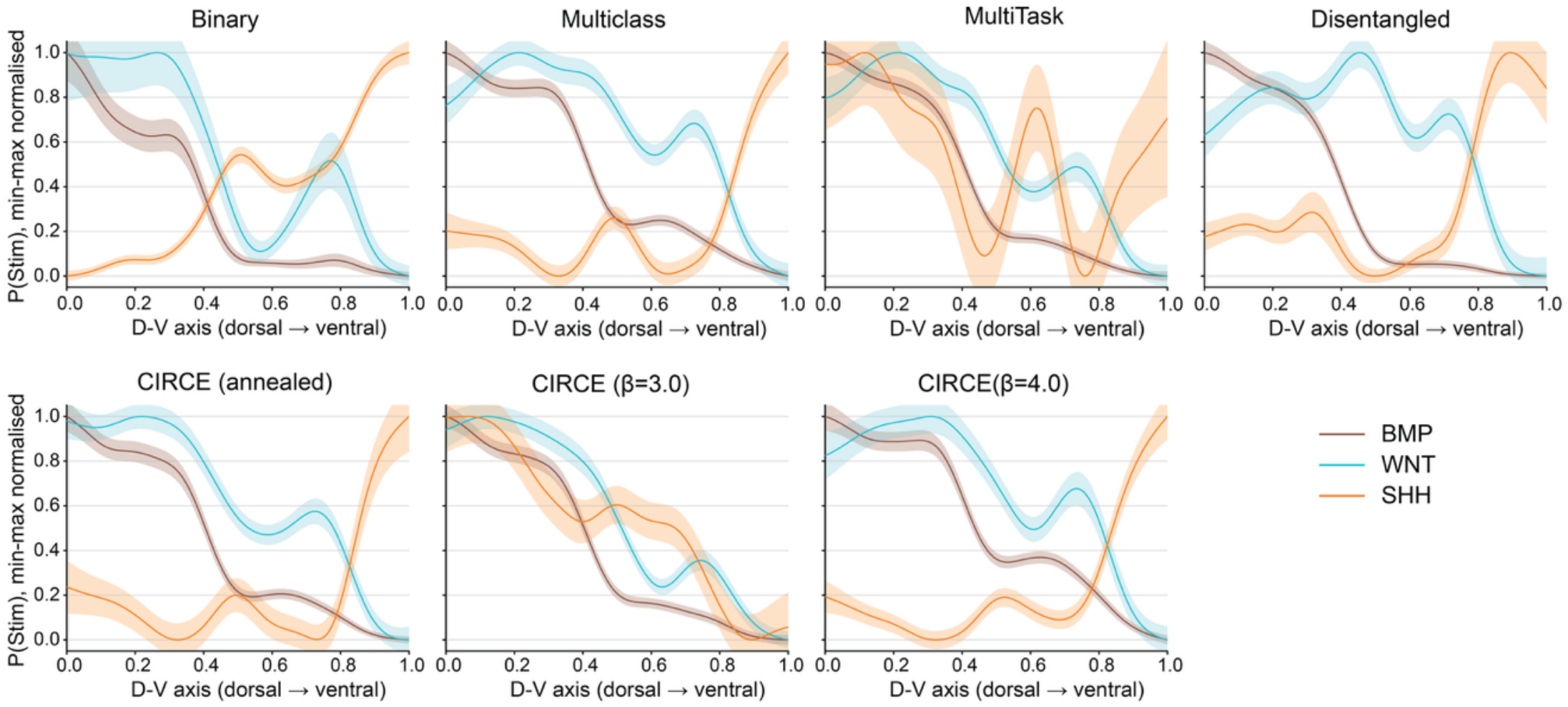
Predicted signaling gradients in the developing neural tube. The developmental trajectory was inferred with diffusion pseudotime and marker genes for the dorsal-ventral axis. Inferred signaling activation probabilities were plotted as mean (solid line) and SEM (shaded area) within each spatial bin along the DV axis. CIRCE extends continuous signaling profiles across the inferred developmental axis relative to models trained with hard labels.

**Supplemental Figure 15.**
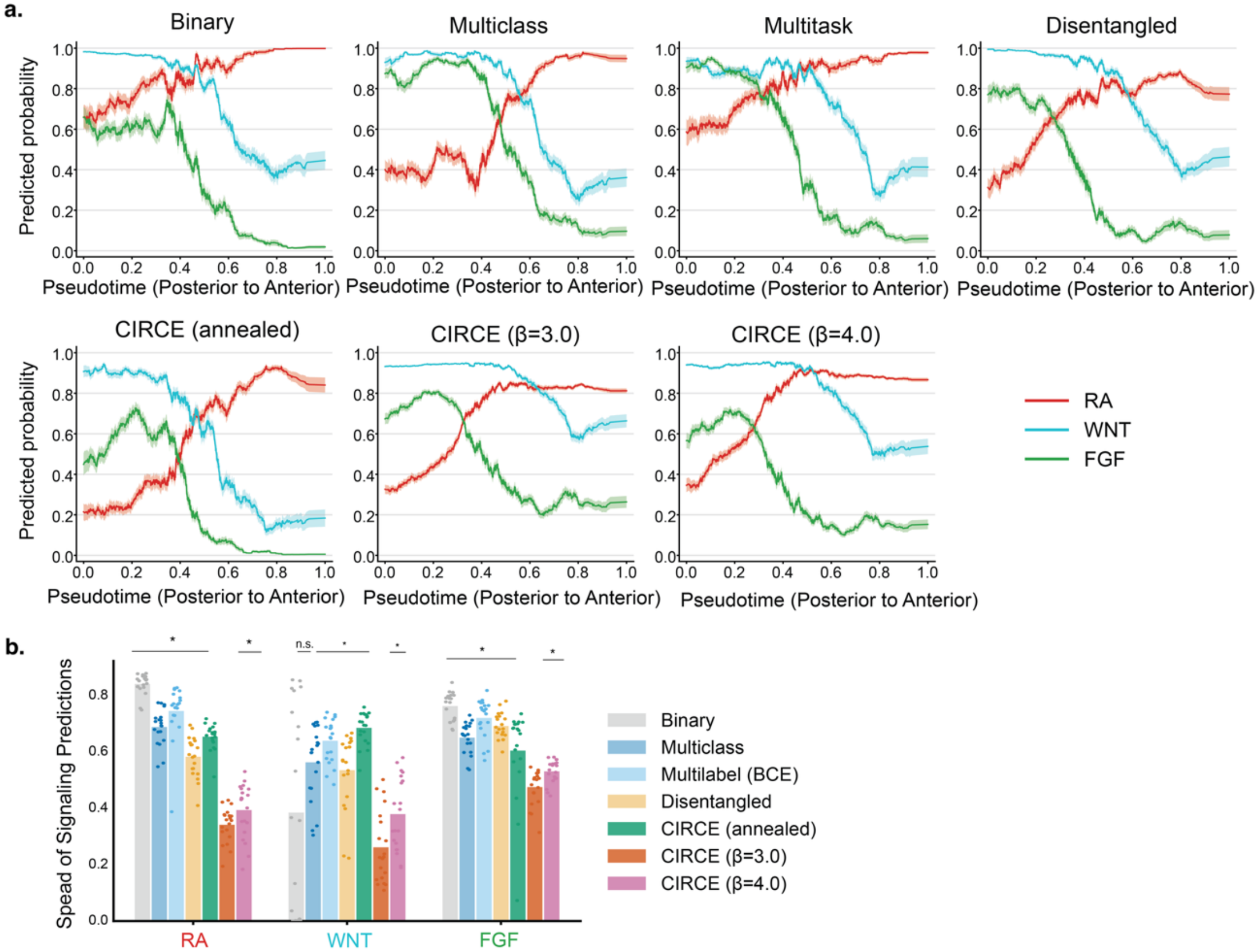
Graded signaling predictions across embryonic somitic mesoderm. **a**, Predicted signaling gradients along the anterior-posterior (AP) axis for the embryonic somitic mesoderm. The developmental trajectory was inferred with diffusion pseudotime and marker genes. Inferred signaling activation probabilities were plotted as mean (*solid line*) and SEM (*shaded area*) within each spatial bin along the AP axis. **b**, Quantifying the spread of predictions. The predicted signal activation probabilities within each spatial bin in pseudo-space are tighter for CIRCE (*β* = 3.0,4.0) models than all other models, comparable to the neural tube results. * denotes p < 0.05 by a two-tailed Wilcoxon ranked signed test with a Benjamini-Hochberg correction for false discovery.

